# PGC-1α drives small cell neuroendocrine cancer progression towards an ASCL1-expressing subtype with increased mitochondrial capacity

**DOI:** 10.1101/2024.04.09.588489

**Authors:** Grigor Varuzhanyan, Chia-Chun Chen, Jack Freeland, Tian He, Wendy Tran, Kai Song, Liang Wang, Donghui Cheng, Shili Xu, Gabriella A. Dibernardo, Favour N Esedebe, Vipul Bhatia, Mingqi Han, Evan R. Abt, Jung Wook Park, Sanaz Memarzadeh, David Shackelford, John K. Lee, Thomas Graeber, Orian Shirihai, Owen Witte

## Abstract

Adenocarcinomas from multiple tissues can evolve into lethal, treatment-resistant small cell neuroendocrine (SCN) cancers comprising multiple subtypes with poorly defined metabolic characteristics. The role of metabolism in directly driving subtype determination remains unclear. Through bioinformatics analyses of thousands of patient tumors, we identified enhanced PGC-1α—a potent regulator of oxidative phosphorylation (OXPHOS)—in various SCN cancers (SCNCs), closely linked with neuroendocrine differentiation. In a patient-derived prostate tissue SCNC transformation system, the ASCL1-expressing neuroendocrine subtype showed elevated PGC-1α expression and increased OXPHOS activity. Inhibition of PGC-1α and OXPHOS reduced the proliferation of SCN lung and prostate cancer cell lines and blocked SCN prostate tumor formation. Conversely, enhancing PGC- 1α and OXPHOS, validated by small-animal Positron Emission Tomography mitochondrial imaging, tripled the SCN prostate tumor formation rate and promoted commitment to the ASCL1 lineage. These results establish PGC-1α as a driver of SCNC progression and subtype determination, highlighting novel metabolic vulnerabilities in SCNCs across different tissues.

**STATEMENT OF SIGNIFICANCE:** Our study provides functional evidence that metabolic reprogramming can directly impact cancer phenotypes and establishes PGC-1α-induced mitochondrial metabolism as a driver of SCNC progression and lineage determination. These mechanistic insights reveal common metabolic vulnerabilities across SCNCs originating from multiple tissues, opening new avenues for pan-SCN cancer therapeutic strategies.

## INTRODUCTION

Epithelial cancers from various tissues, including the prostate, lung, and bladder, can converge to an aggressive small cell neuroendocrine (SCN) phenotype under targeted treatments^1–3^. These phenotypically similar, small cell neuroendocrine cancers (SCNCs) are classified as high-grade, poorly differentiated small cell carcinomas that commonly express neuroendocrine markers like chromogranin A (CHGA), neural cell adhesion molecule 1 (NCAM1), and synaptophysin (SYP)^4^. Small cell lung cancer (SCLC), the most prevalent type of SCNC, represents about 14% of all lung cancer diagnoses, with approximately 200,000 patients succumbing to the disease annually. The prevalence of treatment-induced SCN prostate cancer, which accounts for approximately 10-40% of all androgen deprivation therapy (ADT)-insensitive, castration-resistant prostate cancer (CRPC) cases, is increasing, likely due to resistance mechanisms ADT. With a limited long-term effectiveness of current chemotherapy strategies^5^ indicated by a dismal median survival rate between 7-16 months^6^, there is a pressing need for a better mechanistic understanding of disease progression to inform new therapeutic strategies.

SCNCs were traditionally seen as a uniform group of cancers but are now understood to include multiple subtypes. These subtypes, defined by their prominent expression of lineage-specific molecular markers, showcase the diversity within SCNCs. This is best exemplified in SCLC, which is categorized into ASCL1, NEUROD1, POU2F3, and YAP1 subtypes^7,8^. Recent studies have identified corresponding subtypes in SCN prostate cancer, including ASCL1, POU2F3/ASCL2, and NEUROD1^9–11^. A growing body of research across numerous SCNC models has highlighted the plastic nature of differentiation towards and between these subtypes^7,8,12–14^, a feature that likely contributes to treatment resistance.

Complementing the molecular heterogeneity within SCNCs, recent studies have begun to highlight the metabolic diversity across different SCLC subtypes^15–19^. Further, a correlation has been observed between ASCL1 expression levels and a dependence on inosine monophosphate dehydrogenase, a key enzyme in guanosine biosynthesis critical for normal metabolic functions^15^. These insights raise the possibility that metabolic reprogramming might regulate lineage plasticity in SCNCs. Indeed, metabolic reprogramming not only influences cell fate decisions^20^ but is also orchestrated by oncogenes to regulate these decisions^21^. These observations highlight the intimate interplay between cancer metabolism and lineage plasticity. SCN-associated oncogenes such as MYC^22–26^, AKT^27,28^, TP53^29,30^, and RB1^31–33^, regulate metabolic pathways individually, but their collective impact on metabolic reprogramming and lineage determination during SCNC remains unclear.

A key aspect of cancer cell metabolic reprogramming is the shift towards aerobic glycolysis (also known as the Warburg effect) characterized by an increased rate of glucose consumption and lactate secretion, even in the presence of ample oxygen^34–36^. This seminal discovery made nearly a century ago by Otto Warburg led to a longstanding belief that cancer cells have dysfunctional mitochondria. However, recent studies have countered this perspective, showing that many cancers actually have functional mitochondria fully capable of processing fuels via the tricarboxylic acid (TCA) cycle to support oxidative phosphorylation (OXPHOS)^37^. In fact, some cancers have a heightened reliance on OXPHOS^38,39^, which is sometimes linked with therapy resistance^40–45^. This emerging understanding has sparked significant interest in OXPHOS inhibitors as promising candidates for cancer therapy^46–48^. However, their utility in treating SCNCs remains unclear owing to an incomplete understanding of the metabolic dynamics in these aggressive cancers. Although increased aerobic glycolysis has been observed in SCN prostate cancer^49^, the OXPHOS status in SCNCs and their subtypes remains unclear. Therefore, we reasoned that exploring mitochondrial function in SCNCs may reveal pivotal insights into their pathophysiology

To explore the influence of mitochondrial function on SCNC development, we focused on peroxisome proliferator-activated receptor gamma coactivator 1-alpha (PGC-1α)^50^, a transcriptional co-activator widely recognized as a potent regulator of mitochondrial biogenesis and OXPHOS. Originally identified for its role in cold-induced adaptive thermogenesis through the regulation of OXPHOS genes^51^, PGC-1α has since been implicated in metabolic reprogramming during cancer development, demonstrating robust but varied effects across cancer types^52^. In prostate cancer, it has been found to support the growth of localized, androgen-dependent tumors^53,54^ while restraining the progression of androgen-independent, metastatic forms^55–57^. In melanoma, PGC-1α is associated with tumors that exhibit enhanced mitochondrial function^58^ and a reduced metastatic potential^59^. It suppresses metastasis in non-small cell lung cancer (NSCLC)^60^, but promotes metastasis in other cancer models^61^. These observations demonstrate that while the exploration of PGC-1α has yielded critical insights into cancer metabolism and progression, its influence is context-dependent. Therefore, unraveling the specific effects of PGC-1α in SCNCs could provide key insights into potential therapeutic targets or prognostic factors.

To investigate the role of PGC-1α in regulating SCNC development, we performed an integrated computational and functional investigation. Bioinformatics analyses of 10,529 patient tumors and 1,466 human cancer cell lines were performed, followed by functional and mechanistic analyses using multiple model systems including a human prostate tissue-derived SCN transformation system, referred to as PARCB^2,3,11^. This system enables the transformation of normal human lung epithelial cells, prostate epithelial cells, or bladder urothelial cells into SCNC via transduction with an oncogene cocktail abbreviated as PARCB (T<u>P</u>53DN, myr-<u>A</u>KT, sh<u>R</u>B1, <u>c</u>-MYC, and <u>B</u>CL2). This model generates two distinct tumor lineages characterized by either stem-like (POU2F3/ASCL2 subtype) or neuroendocrine features (ASCL1 subtype) that closely resemble the POU2F3 and ASCL1 subtypes seen in clinical SCNCs^11^. Here, we reveal the metabolic heterogeneity among SCNC subtypes, defined by PGC-1α and OXPHOS levels. We also identify PGC-1α as a driver in the ASCL1 lineage and uncover OXPHOS as a broad metabolic vulnerability across SCNCs originating from multiple tissues.

## RESULTS

### Elevated PGC-1α in clinical SCNC correlates with increased ASCL1 expression and neuroendocrine differentiation

To investigate the role of mitochondrial metabolism in SCNCs, we started by conducting bioinformatics analyses in a wide range of human cancer cell lines, human primary tumors, and clinical SCN datasets. First, a co-expression analysis was conducted on PGC-1α against the four SCNC lineage markers—ASCL1, POU2F3, NEUROD1, and YAP1—across all 1,466 human cancer cell lines from the Cancer Cell Line Encyclopedia (CCLE). While PGC-1α does not correlate with POU2F3, NEUROD1, or YAP1 (**Figure S1A**), it shows significant positive correlation with the lineage marker ASCL1, a pioneer transcription factor that is essential for neuronal commitment and SCNC progression^62–64^ (**Figure 1A**). Notably, we observed a distinct subpopulation of cell lines with high ASCL1 expression that also have high PGC-1α expression (**Figure 1A, left panel**). We refer to this subpopulation as "PGC-1α/ASCL1-High." To determine the cancer type composition of the PGC-1α/ASCL1-High subpopulation compared to the rest of the samples, we quantified the percentage of cell lines with SCN features in both groups. The PGC-1α/ASCL1-High subpopulation is predominantly comprised of SCNCs (54/68, 79%) (**Figure 1A right panel**), whereas only a small fraction of the rest of the cells (88/1466, 6%) are SCNCs. Consistently, PGC-1α expression is elevated in SCLC and neuroblastoma cell lines (two cancers with SCN differentiation) compared with NSCLC, which typically does not exhibit SCN differentiation (**Figure S1B**). These observations suggest that PGC-1α and ASCL1 are co-upregulated in SCNCs from multiple tissues.

**Figure 1.**
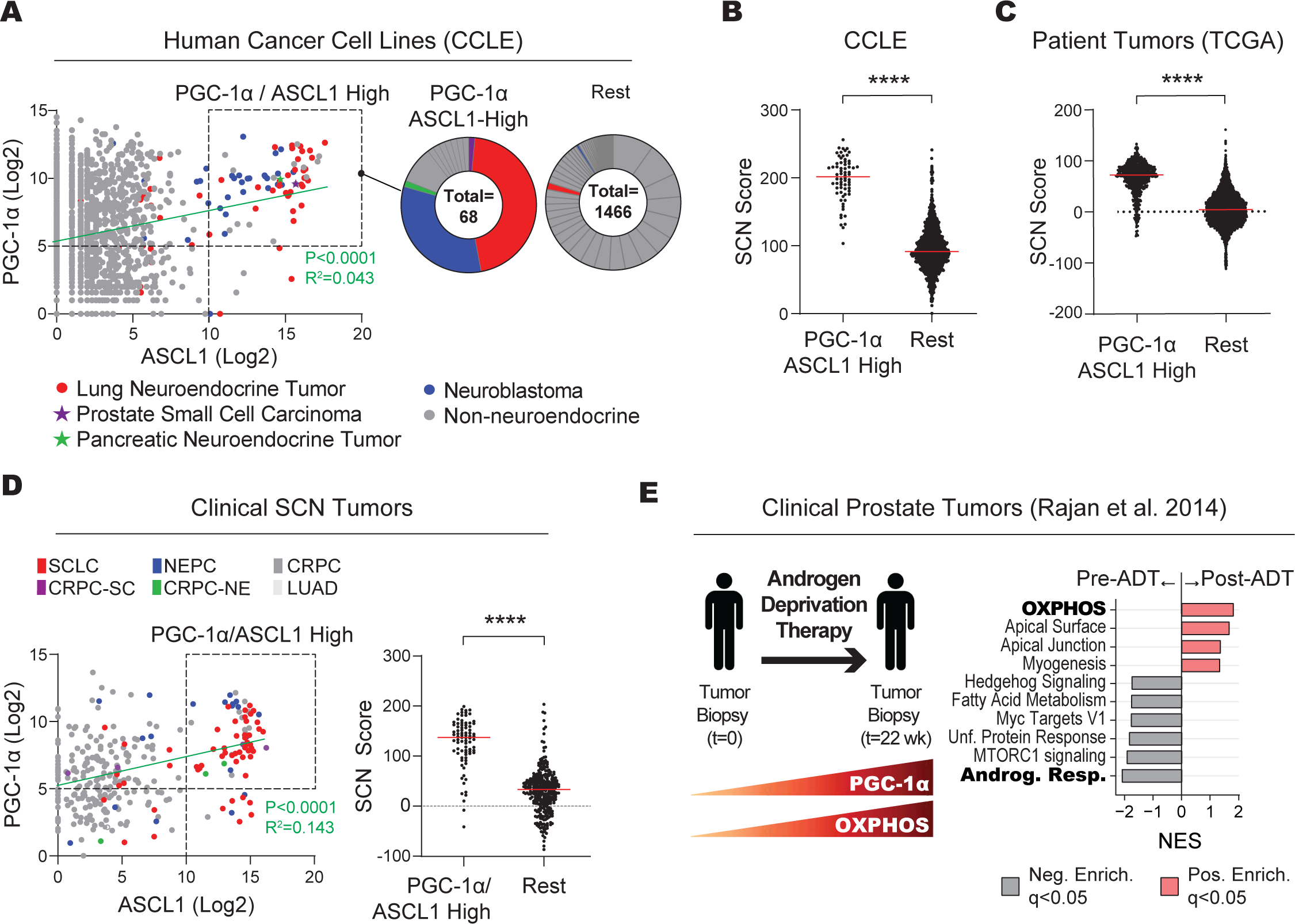
Elevated PGC-1α in clinical SCNC correlates with increased ASCL1 expression and neuroendocrine differentiation. A. Gene expression analysis across all human cancer cell lines from the Cancer Cell Line Encyclopedia (CCLE). Left panel: co-expression analysis between PGC-1α and ASCL1, highlighting samples with high expression levels of both genes (boxed region). Right panel: pie charts illustrating the proportion of cancer types exhibiting elevated co-expression of PGC-1α and ASCL1, in contrast to the rest of the samples. See also Figure S1A. B. Related to Figure 1A and S1B: Transcriptomic examination of small cell neuroendocrine (SCN) differentiation in CCLE human cancer cell lines. SCN scores (defined previously^3^ and detailed in the Methods) are calculated for cells showing heightened PGC-1α and ASCL1 expression versus the rest of the cells. C. Related to Figure S1C: A transcriptomic analysis of SCN differentiation in all the patient tumors from the TCGA database shown in Figure S1C. SCN scores are calculated from the boxed tumor samples with elevated PGC-1α and ASCL1 expression compared with the rest of the samples. D. Gene expression analysis in multiple clinical SCNC datasets^66–70,106^. Left panel: co-expression analysis of PGC-1α and ASCL1. Right panel: SCN scores of the boxed samples with elevated PGC-1α and ASCL1 expression compared with the rest of the samples. E. Transcriptomic analyses in clinical prostate cancer tumors before and after androgen deprivation therapy (ADT) with enzalutamide. Left panel: a schematic overview. Right panel: gene set enrichment analysis (GSEA) of differentially expressed genes in pre-versus post ADT-treated tumors. See also Figure S1F. Data for Figures 1B-1D are expressed as mean ± SEM, with significance denoted as **** (p≤0.0001), ** (p≤0.01), and * (p≤0.05). The green lines represent the best-fit linear regression model, with the indicated R² and p-values. All Log2 values have been upper quartile normalized with a pseudocount of 1 (Log2 UQN+1). For statistical tests used, see Material and Methods section. Related data can be found in Figure S1. Abbreviations: SCLC, small cell lung cancer; NEPC, neuroendocrine prostate cancer; CRPC, castration resistant prostate cancer; LUAD, lung adenocarcinoma.

To determine whether PGC-1α/ASCL1-High cells show increased SCN differentiation, we applied a well-established SCN score derived from a pan-cancer transcriptomic analysis^1^, where a higher SCN score indicates a more SCN-like phenotype (see the Methods section for more details). PGC-1α/ASCL1-High cells from the CCLE show a marked increase in SCN score compared to the rest of the samples, indicating heightened SCN differentiation. (**Figure 1B**). Together, these data uncover a hitherto uncharacterized link between PGC-1α and ASCL1.

To determine whether the link between PGC-1α, ASCL1, and SCN differentiation extends beyond cultured cell lines to patient tumors, we analyzed 10,529 solid tumors from The Cancer Genome Atlas (TCGA)^65^. Consistent with findings from the cancer cell lines, co-expression analysis identified a distinct subpopulation of PGC-1α/ASCL1-High tumors (**Figure S1C**), which exhibited a robust increase in SCN score (**Figure 1C**). We then expanded the analysis to include clinical SCN samples from various cancers, including lung adenocarcinoma (LUAD), castration-resistant prostate cancer (CRPC), SCN prostate cancer (NEPC), and small cell lung cancer (SCLC)^66–69^. Similar to the patterns observed in the CCLE cell lines and TCGA patient tumors, the majority of clinical samples with increased ASCL1 expression also showed high PGC-1α expression (**Figure 1D, left panel)**. Moreover, most PGC-1α/ASCL1-High patient samples were cancer types that display neuroendocrine or small cell features (56/83; 67%) and had a substantial increase in SCN score (**Figure 1D, right panel**). Consistent with these observations, PGC-1α expression is increased in SCN prostate cancer compared with castration resistant prostate cancer (CRPC)^66^ and in SCLC^70^ compared with NSCLC^65^ (**Figure S1D**). Thus, as in the CCLE cell lines and TCGA tumors, PGC-1α and ASCL1 are co-upregulated in clinical tumors with heightened SCN differentiation. Of note, the increased PGC-1α expression in SCLC is largely attributable to the neuroendocrine subtype expressing ASCL1 (SCLC-A), further substantiating the link between PGC-1α and ASCL1 (**Figure S1D, right panel**).

Epithelial ovarian cancers, particularly high-grade serous ovarian carcinomas (HGSOC), share genomic alterations with neuroendocrine tumors including mutations in TP53, loss of RB1, and amplification of MYC^71^. We therefore examined expression of PGC-1α and its correlation with ASCL1 in a cohort of patients with a diagnosis of HGSOC, drawing on data from our unpublished research. A positive correlation between PGC-1α and ASCL1 was observed in this cohort (**Figure S1E**), highlighting a consistent link across multiple clinical sample types.

Given the known contribution of ADT to SCN prostate cancer progression^72,73^, we next examined the effect of ADT on PGC-1α expression in clinical prostate cancer (**Figure 1E**). A clinical transcriptomic dataset was utilized in which tumors were biopsied from the same prostate cancer patients before and after ADT with enzalutamide, a potent and selective antagonist of the androgen receptor^74^. ADT-treated tumors show increased PGC-1α expression, which positively correlates with ASCL1, consistent with our analyses of CCLE, TCGA, and clinical SCNC datasets (**Figure S1F**). To identify enriched pathways upon ADT treatment, we performed gene set enrichment analysis (GSEA) on patient samples before and after ADT. Consistent with the increased PGC-1α expression, this GSEA revealed OXPHOS as a top enriched pathway in post-ADT tumors (**Figure 1E**). As expected, the Androgen Response pathway is a top downregulated pathway in patient tumors post-ADT (**Figure 1E**). Collectively, these data indicate that PGC-1α expression is upregulated in multiple SCNCs and uncover a previously undescribed link between PGC-1α, ASCL1, and SCN differentiation.

### Enhanced PGC-1α expression and OXPHOS activity in SCN prostate cancer within the ASCL1 tumor subtype

Having established a link between PGC-1α and ASCL1 in multiple clinical SCNCs, we next sought to investigate the functional role of PGC-1α in specific SCNCs subtypes. With this goal in mind, we utilized our human tissue-derived PARCB-induced SCN transformation model^11^. This model produces ASCL1 and POU2F3/ASCL2 subtypes, closely resembling clinical SCLC and SCN prostate cancer subtypes (**Figure 2A**). To determine if PGC-1α is linked to the ASCL1 subtype in the PARCB model, we conducted a co-expression analysis across all PARCB time course samples. This analysis revealed a tight and positive correlation between PGC-1α and ASCL1, with the highest expression of both genes exclusively found in the ASCL1 subtype (**Figure 2B**). Corroborating this observation, PGC-1α expression markedly increased in five out of ten PARCB tumor series derived from distinct donor tissue series (**Figure 2C**). Moreover, across all the patient series, the ASCL1 neuroendocrine subtype demonstrated a 16-fold higher expression of PGC-1α compared with the POU2F3/ASCL2 subtype (**Figure 2D)**. To check whether PGC-1α and ASCL1 are co-expressed at the single cell level, we examined single-cell RNA-sequencing (scRNA-seq) from the same PARCB time course dataset^11^. The analysis revealed that high PGC-1α expression was almost exclusively associated with ASCL1, while it was absent in cells expressing POU2F3 or ASCL2 (**Figure 2E**). Collectively, these data indicate increased PGC-1α expression in the ASCL1 subtype, which aligns with the bioinformatics analyses in patient tumors and clinical datasets described in Figure 1.

**Figure 2.**
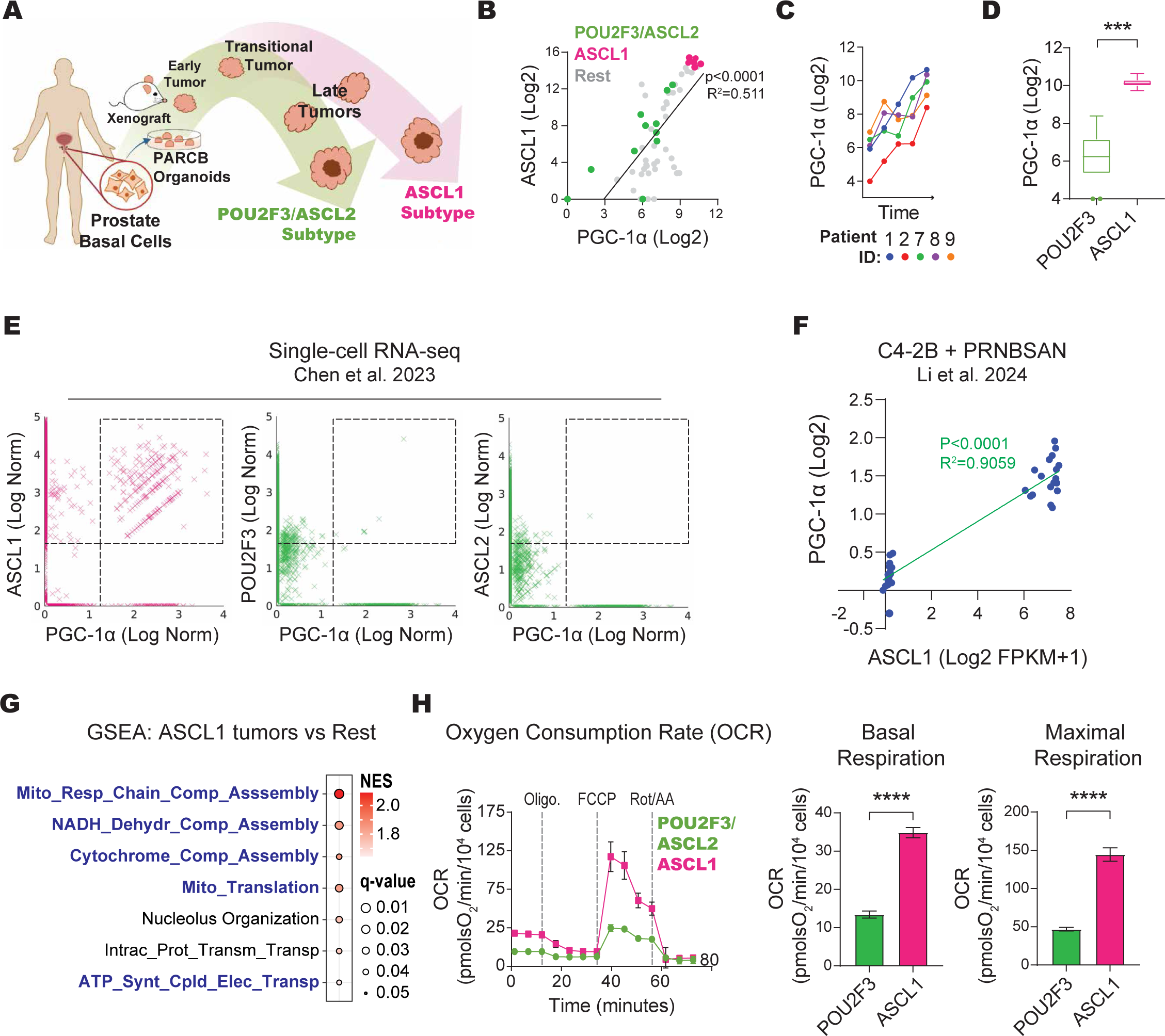
Enhanced PGC-1α expression and OXPHOS activity in SCN prostate cancer within the ASCL1 tumor subtype. A. Schematic illustrating PARCB-induced SCN prostate transformation, highlighting the bifurcation of end stage tumors into the POU2F3/ASCL2 and ASCL1 lineages. The schematic is modified from Chen et al. 2023^11^. B. Co-expression analysis of PGC-1α and ASCL1 during PARCB prostate transformation. Note that tumors with the highest expression levels of both genes are exclusively found in the ASCL1 subtype. The black line represents the best-fit linear regression model, with the indicated R² and p-values. C. Analysis of PGC-1α expression levels during PARCB transformation in 5 out of 10 patient series showcasing increased PGC-1α expression over time. D. Comparative expression levels of PGC-1α in POU2F3/ASCL2 versus ASCL1 PARCB tumor subtypes indicating increased PGC-1α expression in the ASCL1 subtype. E. Single cell gene expression levels of PGC-1α vs ASCL1, POU2F3, and ASCL2 from the PARCB time course tumors from Chen et al. 2023^11^. F. Co-expression analysis of PGC-1α and ASCL1 in C4-2B cells from the PRNBSAN transformation system^77^. Log2 values are Log2 FPKM+1. G. GSEA in the ASCL1 tumor subtype versus the rest of the PARCB time course samples. Gene Ontology Biological Process (GOBP) gene sets were used. The pie chart indicates the proportion of enriched gene sets related to mitochondrial OXPHOS. H. Seahorse respirometry in PARCB tumor-derived cell lines from both the POU2F3/ASCL2 and ASCL1 subtypes. Inhibitors used: Oligo, Oligomycin; FCCP, Carbonyl cyanide-p-trifluoromethoxy-phenylhydrazone; Rot/AA, rotenone / antimycin A. All data, except in Figure 2F, utilizes data extracted from our recently published Chen et al. 2023 dataset^11^. Data in Figures 2D, 2F, and 2H are presented as mean ± SEM, with significance levels marked as follows: **** (p≤0.0001), *** (p≤0.001), ** (p≤0.01), and * (p≤0.05). All Log2 values are Log2 UQN+1 unless otherwise noted. For statistical tests used, see Material and Methods section. Related data can be found in Figure S2-S6.

Building on our findings that PGC-1α expression is elevated in the ASCL1 tumor subtype, we further investigated the regulatory mechanisms underlying this phenomenon. Specifically, we aimed to determine if transcription factors (TFs) co-activated by PGC-1α are more accessible in the ASCL1 subtype than in the POU2F3/ASCL2 subtype. Hypergeometric Optimization of Motif EnRichment (HOMER) analysis was performed to compare the accessible peaks in both tumor subtypes. In the ASCL1 tumor subtype, there was a notable enrichment of transcription factors co-activated by PGC-1α, including HNF4 and PPPARA, which were among the most significantly enriched (**Figure S2**). This enrichment suggests enhanced PGC-1α-induced TF signaling in the ASCL1 subtype, further corroborating the link between PGC-1α and ASCL1.

Given the shared molecular alterations driven by a common set of oncogenes in both ASCL1 and POU2F3/ASCL2 subtypes, we investigated whether ASCL1, as a pioneer transcriptional activator, could directly interact with PGC-1α gene promoters to enhance its expression specifically in the ASCL1 subtype. We analyzed ASCL1 ChIP-Seq datasets from SCN lung^75,76^ and prostate cancer cell lines^63,77^, as well as in LuCaP prostate cancer patient-derived xenografts (PDXs)^10^. ASCL1’s binding to an alternative promoter of PGC-1α, particularly at exon 1b^78^ encoding transcript variant X2 (XM_005248132.1), was modest in SCN lung and prostate cancer cell lines but absent in prostate cancer PDX models (**Figure S3**). In contrast, prostate cancer PDX models showed ASCL1 with a stronger affinity for variant 1 (NM_001330751.2) of PGC-1α, a pattern not observed in SCN lung and prostate cancer cell lines (**Figure S4**). Additionally, ASCL1’s interaction within the PGC-1α gene body was exclusive to SCN lung cancer cell lines and was not present in SCN prostate cancer cell lines and PDX models (**Figure S5**). These inconsistent binding patterns across datasets and models hinder definitive conclusions about ASCL1’s direct regulation of PGC-1α.

Given our findings of variable ASCL1 binding to PGC-1α DNA, we sought to determine if a functional connection exists that does not solely rely on direct DNA interaction. To this end, we investigated whether exogenous ASCL1 expression could upregulate PGC-1α. Analyzing data from our previous study^11^, we found that ASCL1 overexpression in two PARCB POU2F3/ASCL2 tumor-derived cell lines, which naturally do not express PGC-1α, led to increased PGC-1α expression in both cell lines (**Figure S6A**). This enhancement of PGC-1α expression by ASCL1 suggests that ASCL1 can regulate PGC-1α expression even in cells that naturally lack it.

Given our findings of variable ASCL1 binding to PGC-1α DNA, we sought to determine if a functional connection exists that does not solely rely on direct DNA interaction. TO this end we examined whether exogenous expression of ASCL1 could upregulate PGC-1α. We analyzed data from our prior study where ASCL1 was overexpressed in two PARCB POU2F3/ASCL2 tumor-derived cell lines, which do not express PGC-1α^11^ (**Figure S6A**). ASCL1 overexpression enhanced the expression of PGC-1α in both cell lines (**Figure S6A**), indicating that it can regulate PGC-1α expression.

To ensure that ASCL1’s regulatory effect on PGC-1α extends beyond the PARCB cell lines, we utilized a newly developed in vitro model for SCN prostate cancer differentiation using the androgen-insensitive C4-2B cell line^77^. This model induces SCN differentiation by incorporating a specific set of SCN oncogenes and transcription factors, namely PRNBSAN (dominant-negative T<u>P</u>53, sh<u>R</u>B1, MYC<u>N</u>, <u>B</u>CL2, <u>S</u>RRM4 combined with <u>A</u>SCL1, or <u>N</u>EUROD1) (**Figure S6B**). Consistent with findings in the PARCB cell lines, we observed a strong correlation between PGC-1α and ASCL1 expression, evidenced by an R^2^ value of 0.9059 (**Figure 2F**). Moreover, PGC-1α levels correlated with enhanced SCN differentiation in this model (**Figure S6C**). Collectively, these findings across multiple model systems establish ASCL1 as a positive regulator of PGC-1α expression and reinforce the functional connection between PGC-1α, ASCL1, and neuroendocrine differentiation.

Following our demonstration of both a tight correlative and functional link between ASCL1 and PGC-1α, we proceeded to define the transcriptional changes associated with the development of the ASCL1 subtype. GSEA was performed in tumors of the ASCL1 subtype versus all the other stages of PARCB transformation (**Figure 2G**). Consistent with their high PGC-1α expression, ASCL1 tumors had pronounced upregulation of oxidative phosphorylation (OXPHOS) genes (**Figure 2G**), indicating increased mitochondrial biogenesis and/or activity. To determine whether the increased OXPHOS expression translated to higher OXPHOS activity, we performed respirometry using a Seahorse XF Analyzer (Agilent Technologies). This extracellular flux analysis utilizes multiple inhibitors targeting different components of the OXPHOS machinery, providing a detailed functional profile of OXPHOS activity^79^. Seahorse respirometry revealed that PARCB-ASCL1 tumor-derived cell lines exhibited a marked increase in basal and maximal oxygen consumption rates (**Figure 2H**), indicating increased OXPHOS activity compared with the POU2F3/ASCL2 subtype. Furthermore, the ASCL1 tumor subtype had higher expression of both subunits of the mitochondrial pyruvate carrier (MPC) (**Figure S6D**), an obligate heterodimer in the mitochondrial inner membrane that allows entry of pyruvate into the mitochondrion to fuel OXPHOS^80^. Collectively, these findings demonstrate that the ASCL1 subtype exhibits enhanced PGC-1α expression and OXPHOS activity.

### PGC-1α inhibition blunts OXPHOS, reduces the growth of SCNC cell lines, and blocks SCN prostate tumor formation

Considering the increased PGC-1α expression in SCNCs described in Figures 1 and 2, we next asked whether PGC-1α is important for PARCB tumor progression. PGC-1α was inhibited during PARCB prostate transformation using shRNA interference (**Figure S7A**). PGC-1α inhibition was verified using RT-qPCR in PARCB organoids before xenografting into immunocompromised mice for tumor formation (**Figure S7B**). PGC-1α inhibition blocked PARCB tumor growth, underscoring its requirement for PARCB prostate tumor initiation.

Since PARCB tumors could not form under PGC-1α inhibition, we explored its role in the proliferation of PARCB tumor derived prostate cancer cell lines. PGC-1α inhibition reduced the proliferation of both ASCL1 and POU2F3/ASCL2 subtypes with a more pronounced effect in the latter (**Figures 3A**). Next, we examined whether pharmacological inhibition of PGC-1α similarly affects cell proliferation. A recent high-throughput screen identified a small molecule inhibitor called SR-18292 that increases PGC-1α lysine acetylation and inhibits PGC-1α activity by blocking its interaction with HNF4A^81^. As with genetic inhibition, SR-18292 treatment suppressed growth in both ASCL1 and POU2F3/ASCL2 subtypes, with a stronger effect on the POU2F3ASCL2 subtype (**Figure 3B**).

**Figure 3.**
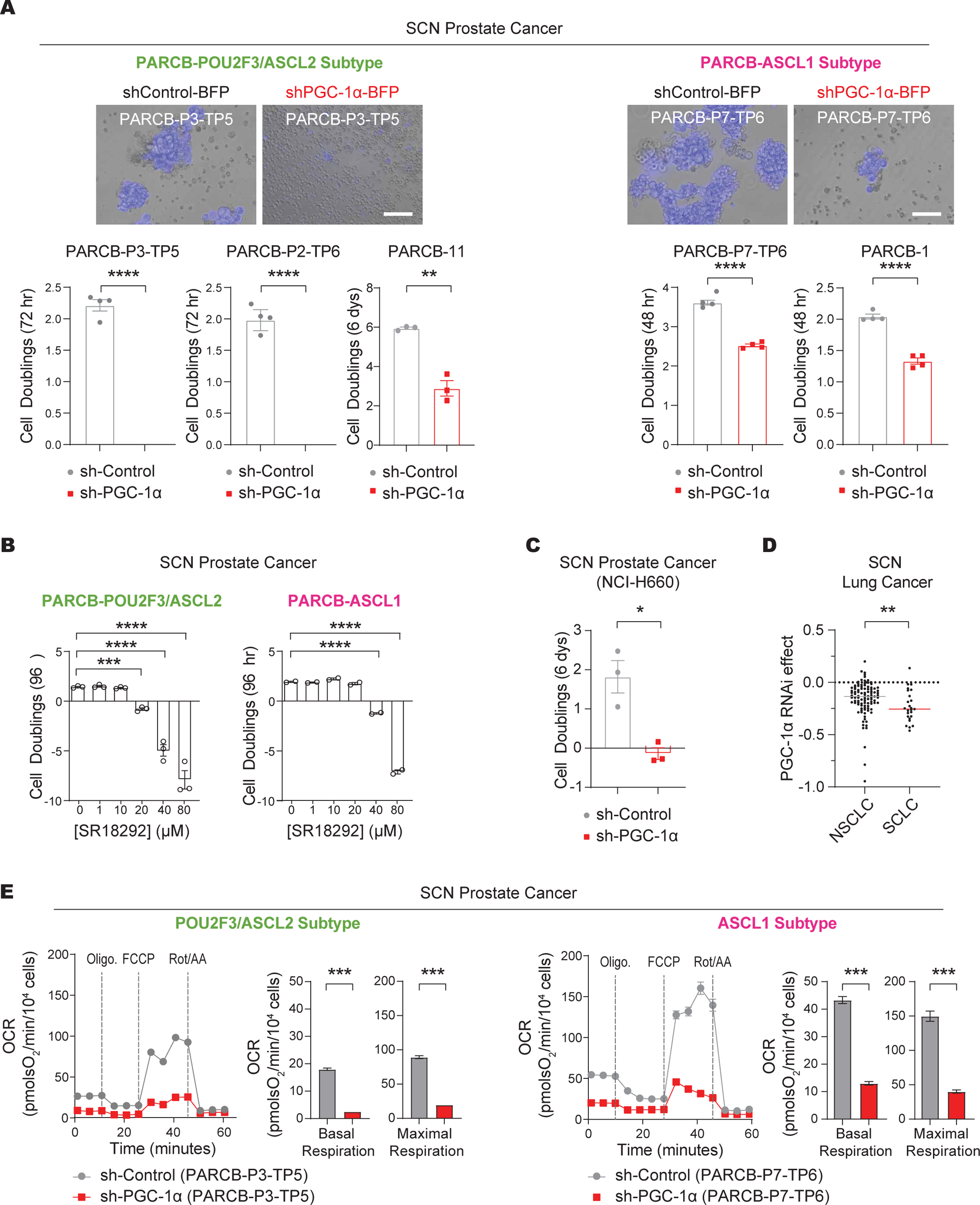
PGC-1α inhibition blunts OXPHOS, reduces the growth of SCNC cell lines, and blocks SCN prostate tumor formation. A. Cell proliferation analysis in PARCB tumor-derived cell lines comparing PGC-1α knockdown (KD) cells to control. The blue fluorescence in the micrographs is derived from the shPGC-1α and shControl constructs, which have co-linked expression of a blue florescence protein (see methods section for details). B. Cell proliferation analysis of PARCB tumor-derived cell lines treated with SR-18292, a pharmacological inhibitor of PGC-1α. C. Cell proliferation analysis of a patient-derived SCN prostate cancer cell line (NCI-H660) with PGC-1α KD compared with control. D. PGC-1α inhibition (RNAi) effects on cell proliferation, represented by a dependency scores from the DepMap database. A score of zero signifies no proliferation impact, negative values indicate decreased proliferation, and a score of negative one corresponds to the median effect across all pan-essential genes. E. Seahorse respirometry in PARCB cell lines derived from the POU2F3/ASCL2 and ASCL1 tumor subtypes. Oligo, Oligomycin; FCCP, Carbonyl cyanide-p-trifluoromethoxy-phenylhydrazone; Rot/AA, rotenone / antimycin A. All data are presented as mean ± SEM, with significance levels marked as follows: *** (p≤0.001), ** (p≤0.01), and * (p≤0.05). For statistical tests used, see Material and Methods section. Related data can be found in Figure S7.

To determine whether this effect of PGC-1α inhibition extends beyond the PARCB-derived cell lines, we first examined its effect on PRNBSA-induced SCN prostate cancer progression (model described in Figure 2C). PGC-1α inhibition using shRNA, blocked the PRNBSA-mediated growth of C4-2B and LNCaP cell lines (**Figure S7C and S7D**). Next, we tested the effect of PGC-1α inhibition on patient-derived SCN prostate and lung cancer cell lines. PGC-1α inhibition reduced the growth of the sole patient-derived SCN prostate cancer cell line, NCI-H660 (**Figure 3C**). Similarly, analysis of PGC-1α inhibition in the Cancer Dependency Map (DepMap)^82^ revealed a stronger anti-proliferative effect in SCLC compared with NSCLC (**Figure 3D**). Taken together, these findings indicate that PGC-1α inhibition blunts the growth of both SCN prostate and lung cancer cell lines.

We next sought to understand the functional mechanism behind PGC-1α inhibition’s impact on SCNC cell line growth and PARCB tumor suppression. We noted in Figure 2 above that the ASCL1 subtype has enhanced PGC-1α expression and OXPHOS activity. We therefore examined whether PGC-1α inhibition would blunt OXPHOS in SCN prostate cancer cell lines. Accordingly, we inhibited PGC-1α in ASCL1 and POU2F3/ASCL2 cell lines using shRNA and performed Seahorse respirometry to examine the effect on mitochondrial function. PGC-1α inhibition downregulated basal and maximal respiration in both the ASCL1 and POU2F3/ASCL2 subtypes (**Figure 3E**), highlighting the importance of PGC-1α in maintaining OXPHOS activity across both subtypes. These findings collectively suggest that PGC-1α is essential for the initiation of PARCB prostate tumors and sustaining OXPHOS and proliferation of cultured SCNC cell lines.

### OXPHOS inhibition blunts SCN prostate and lung cancer cell line proliferation

Because PGC-1α inhibition blunted OXPHOS and reduced the growth of SCN prostate and lung cancer cells, we tested whether OXPHOS is required for the proliferation of SCNC cell lines. First, PARCB cell lines derived from the ASCL1 and POU2F3/ASCL2 tumor subtypes were treated with IMT1B, a recently developed small molecule inhibitor of mitochondrial RNA polymerase (POLRMT), which blocks mitochondrial DNA transcription thereby downregulating all of the mitochondrially-encoded respiratory chain complexes^83^. IMT1B treatment blunted cell proliferation in both SCN prostate and lung cancer cell lines with approximately twice the potency in the POU2F3/ASCL2 subtype (**Figure 4A**).

**Figure 4.**
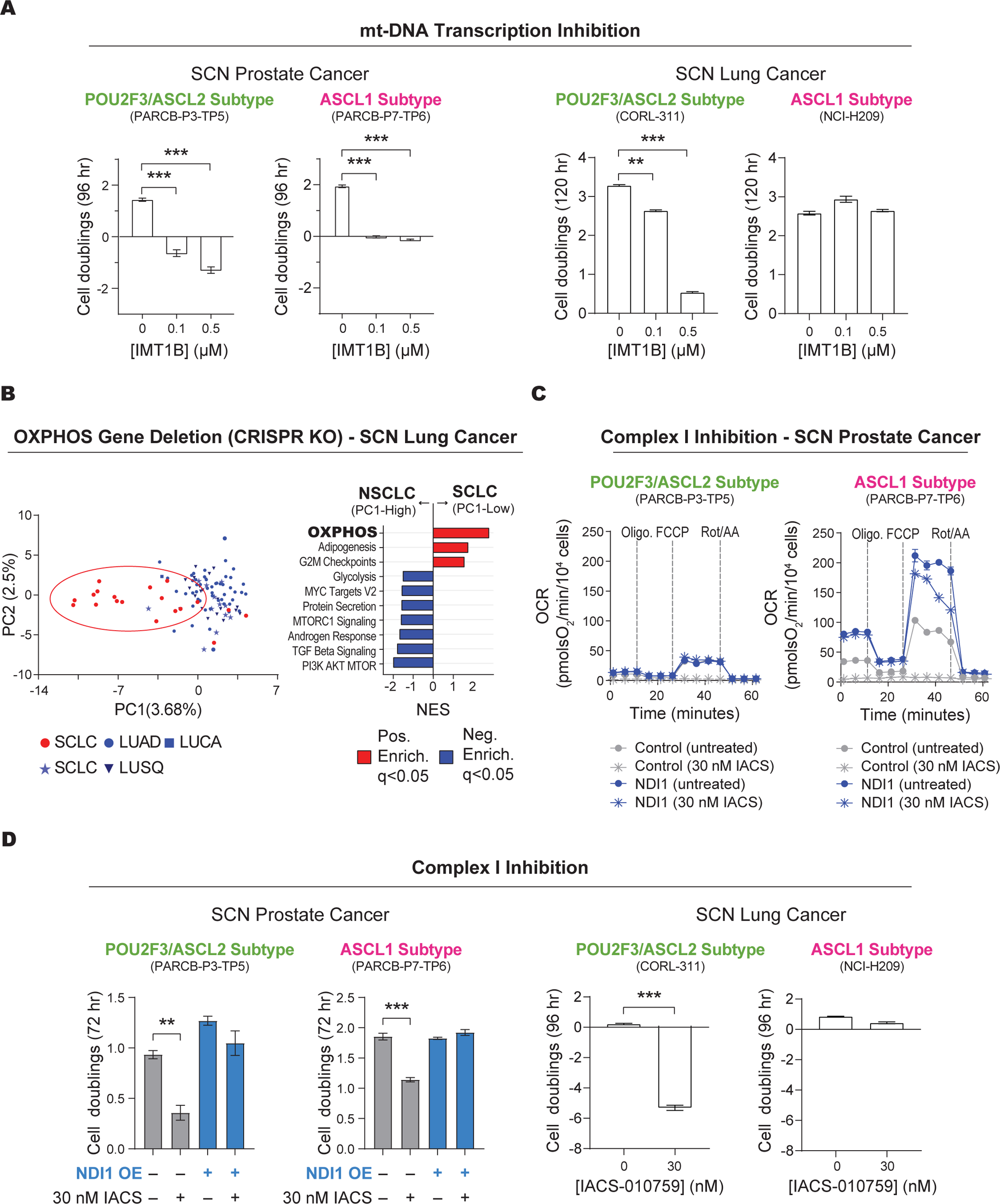
OXPHOS inhibition blunts SCN prostate and lung cancer cell line proliferation. A. Cell proliferation analysis of cell lines derived from the SCN prostate and lung cancer POU2F3/ASCL2 and ASCL1 tumor subtypes treated with IMT1B, a mitochondrial DNA-directed RNA polymerase (POLRMT) inhibitor to block OXPHOS. B. Analysis of differentially dependent genes in NSCLC and SCLC cell lines. Left panel: unsupervised principal component analysis (PCA) on the genetic dependency data from all of the lung cancer cell lines in the DepMap dataset. Right panel: normalized enrichment scores of gene sets that separate PC1 low (SCLC) and PC1 high (NSCLC). The top 100 genes from the PC1 loading scores were used as input. C. Seahorse respirometry in cell lines derived from the PARCB POU2F3/ASCL2 and-ASCL1 tumor subtypes with the indicated conditions. Oligo, Oligomycin; FCCP, Carbonyl cyanide-p-trifluoromethoxy-phenylhydrazone; Rot/AA, rotenone / antimycin A. D. Cell proliferation analysis of cell lines derived from the SCN prostate and lung cancer POU2F3/ASCL2 and ASCL1 subtypes with the indicated treatments. Data in Figure 4B are represented as mean ± SEM. with statistical significance denoted as *** (p≤0.001) and ** (p≤0.01). For statistical tests used, see Material and Methods section.

We next examined the effect of OXPHOS gene deletion in SCN lung cancer by investigating genetic dependencies of 104 lung cancer cell lines in the DepMap database. We first performed unsupervised principal component analysis (PCA) using the DepMap dependency scores. There was a distinct separation between SCLC and NSCLC samples (**Figure 4B, left panel**), indicating unique genetic dependencies between the two groups. Pathway enrichment analysis from the top and bottom 100 PC1 loading genes indicated that OXPHOS is a top enriched pathway in SCLC versus NSCLC cancer cell lines, implying that OXPHOS gene deletion has a stronger anti-proliferation effect in SCLC versus NSCLC (**Figure 4B, right panel**).

Having established that OXPHOS inhibition, through gene deletion or mt-DNA transcription inhibition with IMT1B, reduces SCNC proliferation, we utilized an alternative approach by employing a clinical-grade respiratory chain complex I inhibitor, IACS-010759. This method offered another avenue to explore OXPHOS suppression in SCN prostate and lung cancer cell lines, under conditions that mimic potential clinical interventions. This treatment abolished OXPHOS in cell lines derived from both subtypes (**Figure 4C**). To confirm the inhibitor’s specificity and ensure that the observed effects were directly due to complex I inhibition, NDI1, a yeast complex I^47,84^ was overexpressed concurrently with complex I inhibition. NDI1 overexpressed fully rescued the OXPHOS inhibition caused by IACS-010759 (**Figure 4C**). Interestingly, NDI1 overexpression boosted the oxygen consumption rate (OCR) above baseline levels only in the ASCL1 subtype (**Figure 4C**), suggesting that this subtype not only has increased OXPHOS activity but may also have higher metabolic plasticity compared to the POU2F3/ASCL2 subtype.

Having verified that IACS-010759 blocks OXPHOS in these SCN prostate cell lines, we next asked whether its treatment would reduce their proliferation in culture. In line with the PGC-1α and OXPHOS inhibition experiments described above, IACS-010759 treatment reduced proliferation of cell lines derived from both subtypes, with a more pronounced effect in the POU2F3/ASCL2 subtype (**Figure 4D**). These anti-proliferation effects caused by IACS-010759 were fully rescued by NDI1 overexpression (**Figure 4D**), indicating high specificity of the inhibitor. A similar anti proliferation effect was observed in SCN lung cancer cell lines, with a more pronounced effect in the POU2F3 subtype (**Figure 4D**). Collectively, these results indicate that PGC-1α and OXPHOS inhibition are critical metabolic vulnerabilities in SCN cancers of both the prostate and lung.

### PGC-1α promotes OXPHOS in the prostate cancer ASCL1 subtype

As described above in Figure 3E, PGC-1α inhibition reduced OXPHOS activity in SCNC cell lines. Could overexpressing PGC-1α, therefore, be sufficient to promote OXPHOS and induce lineage plasticity? To test this, we generated a lentiviral PGC-1α overexpression construct for stable transduction. First, PGC-1α overexpression was verified using western blot analysis in a panel of PARCB prostate cancer cell lines generated previously^2^. As expected, PGC-1α overexpression increased both PGC-1α and OXPHOS protein levels in these cell lines (**Figure S8A**). Next, PGC-1α was overexpressed in POU2F3/ASCL2 and ASCL1 PARCB tumor-derived cell lines, and its effect on OXPHOS activity and neuroendocrine differentiation were measured. This analysis utilized Seahorse respirometry, followed by in vivo micro positron emission tomography (PET) and computed tomography (microPET/CT) imaging, and histological analyses **(Figure 5A)**. PGC-1α overexpression upregulated OXPHOS activity in cell lines derived from the ASCL1 but not POU2F3/ASCL2 subtype (**Figure 5B**). This result is consistent with the ASCL1 subtype’s selective OXPHOS enhancement from yeast Complex I overexpression seen in Figure 4A, suggesting a relatively flexible metabolic state in the ASCL1 subtype.

**Figure 5.**
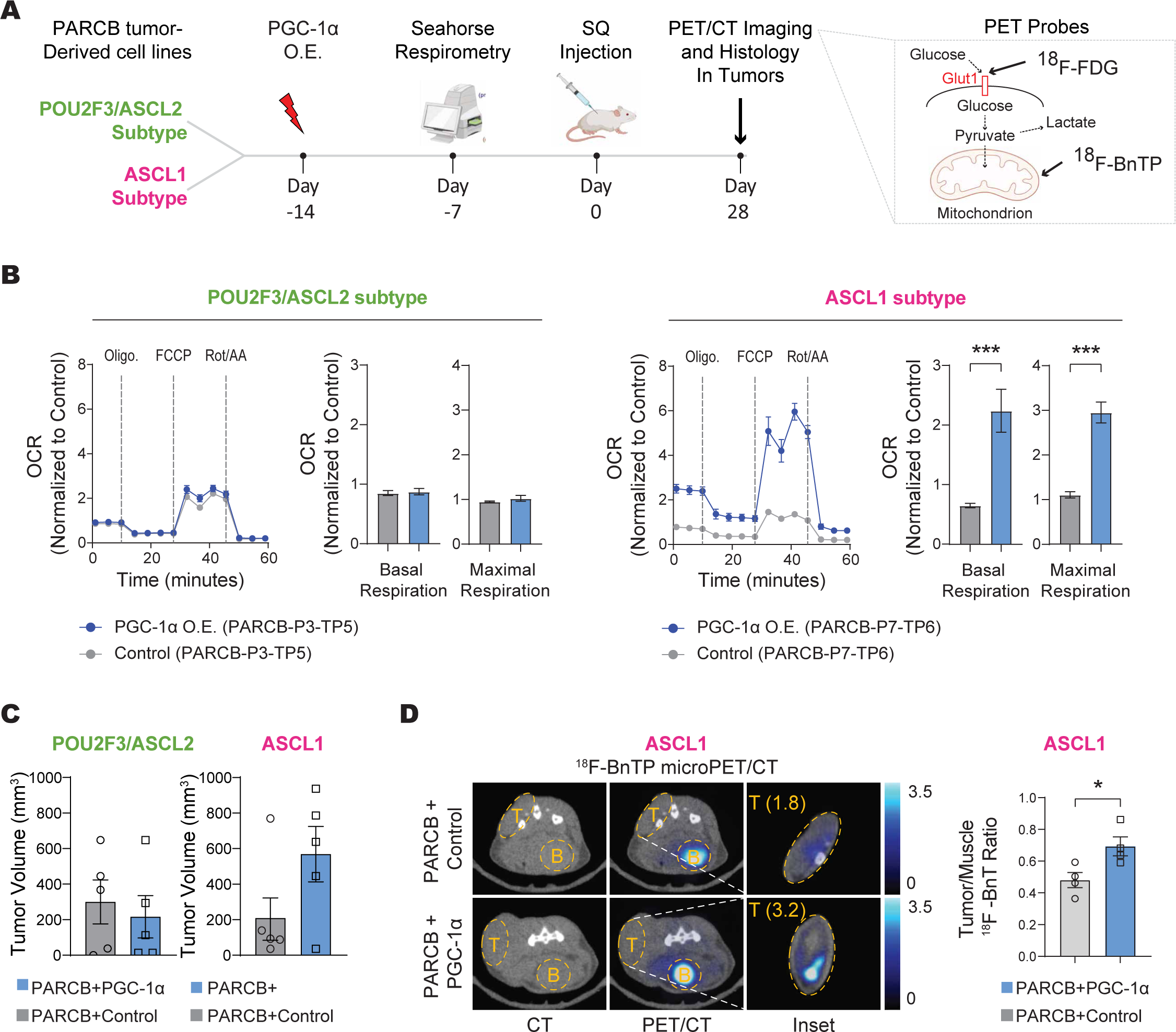
PGC-1α overexpression promotes OXPHOS in the SCN prostate cancer ASCL1 subtype. A. Left panel: overview of experimental strategy. Right panel: Details on the Positron Emission Tomography (PET) probes utilized and their general mechanisms of action. B. Seahorse respirometry in POU2F3/ASCL2 and ASCL1 PARCB tumor-derived cell lines. Oligo, Oligomycin; FCCP, Carbonyl cyanide-p-trifluoromethoxy-phenylhydrazone; Rot/AA, rotenone / antimycin A. C. Caliper measurements of tumors initiated by subcutaneous injection of POU2F3/ASCL2 and ASCL1 PARCB cell lines with PGC-1α overexpression versus control. D. Left panel: Representative ^18^F-FBnTP transverse PET-CT images of mice with subcutaneous tumor implantation. Uptake of PET probe was measured as the maximum percentage of injected dose per cubic centimeter (ID%/cc). Tumors are labeled “T”, and bladders are labeled “B”. The inset has been contrast enhanced to show the relatively low uptake of this probe in subcutaneous tumors, as reported previously^107^. Right panel: quantification of ^18^F-BnTP uptake in the indicated groups. Values are normalized to PET signal from adjacent skeletal muscle. Data are presented as mean ± SEM, with *** indicating a p-value ≤0.001. The absence of asterisks indicates that statistical significance of p≤0.05 was not reached. For statistical tests used, see Material and Methods section. Related data can be found in Figure S8.

To check whether the increased OXPHOS caused by PGC-1α overexpression in cell lines would translate to higher bioenergetics in tumors, ASCL1 or POU2F3/ASCL2 cell lines with and without PGC-1α overexpression were xenografted into immunocompromised mice for tumor initiation and subsequent microPET/CT imaging. PGC-1α overexpression promoted tumor growth only in the ASCL1 subtype, with responses varying from a 2-to 5-fold increase in tumor volume (**Figure 5C**). To evaluate the metabolic changes induced by PGC-1α overexpression in these tumors, we employed two distinct PET probes for microPET/CT imaging, ^18^F-FDG and ^18^F-BnTP (**Figure 5A, S8B, and S8C**). ^18^F-FDG, a glucose analog, is extensively used in clinical PET imaging to measure cellular glucose uptake. While its accumulation in tumors indicates increased glucose uptake, it does not provide information on the specific metabolic pathways, such as whether glucose gets incorporated into the TCA cycle to fuel OXPHOS. To address this limitation, we also utilized ^18^F-BnTP, a lipophilic cation designed to accumulate specifically in the mitochondrial inner membrane and measure mitochondrial membrane potential^85^. This accumulation labels mitochondria with intact OXPHOS activity, offering a direct measure of mitochondrial activity and content in tumors in vivo. PGC-1α overexpression enhanced uptake of both ^18^F-BnTP (**Figure 5D**) and ^18^F-FDG (**Figure S8D**), but only in the ASCL1 subtype. However, the FDG uptake did not reach statistical significance. Collectively, these findings suggest that PGC-1α overexpression promotes tumor growth in the ASCL1 subtype by upregulating mitochondrial activity.

We next checked whether PGC-1α overexpression in tumors derived from ASCL2/POU2F3 or ASCL1 cell lines resulted in increased SCN differentiation. We performed histological analysis from tumors derived from both subtypes. There were no obvious histological changes in SCN differentiation upon PGC-1α overexpression in either subtype (**Figure S8E**). Collectively, the data indicate that, relative to the POU2F3/ASCL2 subtype, the ASCL1 subtype demonstrates dynamic metabolic plasticity, exhibiting a marked OXPHOS response to PGC-1α and NDI1 overexpression. Moreover, the data indicate that despite the ASCL1 subtype’s greater OXPHOS response to PGC-1α overexpression, PGC-1α alone is not sufficient to trigger additional neuroendocrine differentiation in these terminally differentiated cell lines.

### PGC-1α upregulates OXPHOS and drives SCN prostate cancer towards an ASCL1-expressing lineage

We showed in Figure 5 that PGC-1α overexpression was not sufficient to induce lineage plasticity of POU2F3/ASCL2 cells towards ASCL1, nor was it sufficient to induce further neuroendocrine differentiation in terminally differentiated ASCL1 cells. We therefore asked whether PGC-1α overexpression could promote SCN progression during PARCB transformation when the PARCB oncogenes are actively restructuring the chromatin landscape to alter global gene expression. For this, we conducted PARCB prostate transformation experiments with PGC-1α overexpression or control constructs linked with BFP reporters, into the PARCB oncogene cocktail. (**Figure 6A**). PGC-1α overexpression resulted in an approximate 100-fold increase in PGC-1α mRNA levels in PARCB prostate organoids (**Figure S9A**) and a threefold higher tumor take rate compared with controls. (**Figure 6B**). The majority of PGC-1α overexpressing tumors maintained high BFP expression months after transduction, suggesting forward selection during transformation (**Figure S9B**). Consistently, PGC-1α protein levels were higher in PGC-1α overexpressing tumors compared with control (**Figure S9C**). These data indicate that PGC-1α overexpression promotes PARCB-induced SCN prostate tumor formation.

**Figure 6.**
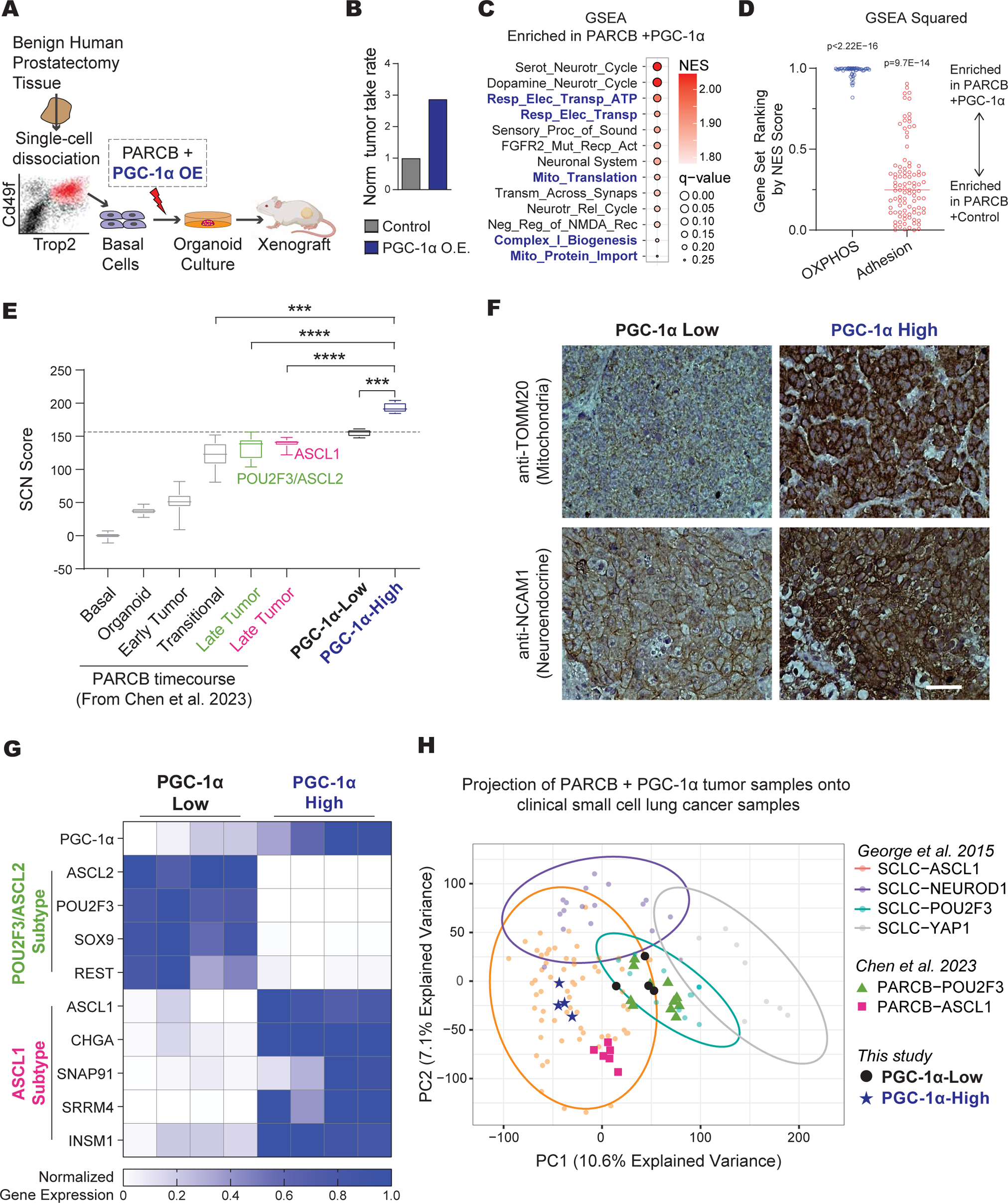
PGC-1α overexpression upregulates OXPHOS and drives SCN prostate cancer towards an ASCL1-expressing lineage. A. Schematic illustrating the experimental setup for assessing the effects of PGC-1α overexpression versus a control vector on PARCB-induced prostate cancer transformation. B. Comparison of the tumor establishment rate (“tumor take rate”) between PARCB transformation using PGC-1α overexpression compared with control. PARCB transformations were performed using three separate patient donors, resulting in a total of 45 xenografts being implanted in 23 mice. C. GSEA in PARCB tumors with PGC-1α overexpression versus control. D. GSEA squared analysis (described in the methods section and in reference^1^) depicting the distribution of normalized enrichment scores (NES) for thousands of gene sets across multiple gene set families in samples with PGC-1α overexpression versus control. E. A transcriptomic analysis of SCN differentiation comparing SCN scores in PGC-1-High versus PGC-1α-Low PARCB tumors. SCN scores (described in the methods section and in reference^1^) calculated from datasets derived from Chen et al 2023^11^ are also shown for comparison. F. Immunohistochemistry in histological sections derived from PARCB tumors with high and low levels of PGC-1α. G. A heatmap displaying normalized gene expression levels for signature genes related to POU2F3/ASCL2 and ASCL1 tumor subtypes in PGC-1α-High versus PGC-1α-Low tumors. H. Principal Component Analysis (PCA) of PARCB samples, distinguishing between PGC-1α-High and PGC-1α-Low PARCB tumors from this study, overlaid on PARCB tumors from our prior time course study^11^ and SCLC cell lines from the CCLE and SCLC patient tumors from George et al. 2015^70^. Data in Figure 6E are presented as mean ± SEM, with statistical significance indicated by **** (p≤0.0001) and *** (p≤0.001). For statistical tests used, see Material and Methods section. Related data be can found in Figure S9.

To better understand how PGC-1α overexpression promotes PARCB transformation, we performed bulk RNA sequencing on PARCB tumors with PGC-1α overexpression and control. GSEA analysis showed that gene sets related to OXPHOS were among the most significantly enriched following PGC-1α overexpression in PARCB tumors (**Figure 6C**). Next, a detailed GSEA squared analysis^1^ (described in the methods section, and in reference^1^) was performed, ranking OXPHOS-related gene sets by NES scores across multiple gene set families comprised over 10,000 gene sets (**Figure 6D**). This analysis showed that OXPHOS gene sets were highly and exclusively enriched in PARCB tumors with PGC-1α overexpression compared with control (**Figure 6D**). Thus, PGC-1α overexpression robustly increases OXPHOS gene expression. Additionally, we explored adhesion-related pathways, known to be broadly downregulated as cancers transition to a more SCN phenotype^1^. We observed a downregulation of adhesion pathways in PARCB tumors that overexpressed PGC-1α, relative to the control group (**Figure 6D**). These finding suggests that PGC-1α overexpression may promote SCN differentiation by upregulating OXPHOS.

To quantify the effect of PGC-1α overexpression on SCN differentiation, we first categorized tumors into PGC-1α-High and PGC-1α-Low groups, based on PGC-1α mRNA levels achieved by overexpression. PGC-1α-high samples exhibited a higher SCN score compared with PGC-1α-low (**Figure 6E**). We also calculated SCN scores for all 55 samples from our recently published PARCB time course study^11^. Notably, PGC-1α-high samples not only displayed increased SCN differentiation in comparison to the PGC-1α-low samples from this study, but also exhibited higher SCN scores than all samples from the temporal study (**Figure 6E**). To determine whether these transcriptomic changes were mirrored at the protein level, we performed immunohistochemistry in PARCB tumors sections with PGC-1α overexpression or control. Consistent with the transcriptomic results in Figures 6C-6E, increased mitochondrial content (TOMM20) and neuroendocrine (NCAM1) expression were observed in PGC-1α-High tumor sections compared with PGC-1α-Low (**Figure 6F**). These results indicate that high levels of PGC-1α overexpression upregulate OXPHOS expression and promote SCN differentiation.

Building on the established relationship between PGC-1α and ASCL1 described above, we investigated if PGC-1α overexpression could promote lineage plasticity towards the ASCL1 subtype, which has high neuroendocrine differentiation. By conducting a gene expression analysis on PARCB tumors categorized by high versus low PGC-1α expression and analyzing signature genes of both ASCL1 and POU2F3/ASCL2 subtypes, we detected a distinct expression pattern. Notably, PGC-1α-high samples predominantly expressed genes associated with the ASCL1 subtype, showing a 14-fold increase in expression compared to PGC-1α-low samples. In contrast, PGC-1α-low samples exhibited nearly a 100-fold increase in the expression of POU2F3/ASCL2 subtype genes compared to PGC-1α-high samples (Figure 6G). (**Figure 6G**). This pattern underscores PGC-1α’s potential influence in steering lineage specificity towards the ASCL1 lineage.

To determine if these subtype-specific expression patterns are observable in a broader clinical context, our datasets from PARCB tumors with PGC-1α overexpression were projected onto datasets from SCLC samples in the CCLE and from the clinical study by George et al. 2015^70^ (**Figure 6H**). We also included the ASCL1 and POU2F3/ASCL2 PARCB samples from our recently published study^11^. PGC-1α-high samples clustered well with SCLC-ASCL1 and PARCB-ASCL1 tumors (HC6), whereas PGC-1α-low samples clustered better with SCLC-POU2F3 and PARCB-POU2F3/ASCL2 tumors (HC5). Collectively, these data suggest that PGC-1α promotes SCNC progression towards an ASCL1-expressing subtype with increased OXPHOS capacity.

## DISCUSSION

### PGC-1α-induced OXPHOS as a driver of lineage plasticity and progression in SCNCs

Metabolic reprogramming is recognized as a hallmark of many cancers, yet its direct role in driving cancer progression and lineage determination remains unclear. Our findings reveal that mitochondrial metabolism, driven by PGC-1α, is not only tightly correlated with SCNC subtypes but also actively drives SCNC development and lineage plasticity toward the ASCL1 subtype, despite similar oncogenic stimuli across subtypes. This is supported by a robust link between PGC-1α and ASCL1 in thousands of human cancer cell lines and patient tumors, as well as a functional connection observed in multiple model systems. These insights pave the way for further investigations into the metabolic underpinnings of SCNCs, suggesting that targeted manipulation of PGC-1α or OXPHOS could offer new avenues for controlling SCNC progression and differentiation.

### ASCL1 as an upstream regulator of PGC-1α

PGC-1α is influenced by a remarkably diverse array of signals, including hormonal, nutritional, and physiological stimuli, as well as potentially hundreds of transcription factors^86^. Despite its complex regulatory network, PGC-1α decisively shapes specific SCNC phenotypes, effectively overriding the oncogenic signals from the same set of oncogenes shared across these subtypes. This observation raises an important question: what specifically regulates PGC-1α in SCNCs in a subtype-specific manner? Our bioinformatics analysis, illustrated in Figure 1, along with functional experiments shown in Figure 2, identify ASCL1 as a regulator of PGC-1α. Moreover, experiments depicted in Figure 6 suggest that PGC-1α fosters lineage plasticity towards the ASCL1 subtype, hinting at a potential co-regulatory mechanism between these two potent transcriptional regulators. While analysis of existing ASCL1 ChIP-Seq datasets identified potential ASCL1 binding sites on PGC-1α promoters and gene body, inconsistencies across these datasets—likely due to variations in experimental conditions and ASCL1 antibodies—render the evidence of direct binding inconclusive. Further research is thus necessary to define the precise interaction between PGC-1α and ASCL1, including exploring indirect pathways by which ASCL1 may influence PGC-1α levels.

### PGC-1α and OXPHOS as metabolic vulnerabilities in SCNC subtypes

OXPHOS inhibition via CRISPR-mediated gene deletion, small molecule inhibitors, or PGC-1α inhibition had an anti-proliferation effect in SCN cancers of both the prostate and lung. These finding have clinical implications given the significant interest in targeting mitochondrial metabolism in various types of cancer^46–48^. The OXPHOS inhibitor IACS-010759, targeting Complex I^46,87–89^, was recently tested in clinical trials. Although initial trials showing peripheral neurotoxicity and lactic acidosis^90^ (on-target, off-tumor toxicities), IACS-010759 demonstrated a partial response and alleviation of cancer-related pain in a patient with advanced castration-resistant prostate cancer^87,91^. While recent clinical trials involving IACS-010759 uncovered significant side effects, SCNCs, particularly of the POU2F3/ASCL2 subtype, may offer a broader therapeutic window for OXPHOS inhibitors. These inhibitors could also be used as co-therapies to enhance their efficacy in SCNCs. Targeting PGC-1α itself is another potential strategy for controlling SCNC progression, with several small-molecule inhibitors that regulate PGC-1α-related pathways^92^. Notably, SR-18292^81^ shown to inhibit PGC-1α^93,94^, reduced proliferation in SCNC-derived cell lines, particularly in the POU2F3/ASCL2 subtype. While this inhibitor was reported to diminish OXPHOS activity^95^, its PGC-1α-related roles beyond OXPHOS in SCNCs warrant further investigation. Collectively, our findings identify the PGC-1α/OXPHOS axis as a novel metabolic vulnerability in SCNCs from multiple tissues.

### The role of PGC-1α and OXPHOS during androgen deprivation therapy

Recent studies have shown that prostate cancer cells increasingly depend on OXPHOS following androgen deprivation therapy (ADT)^96,97^. Our analysis of a clinical dataset, which included tumors sampled from prostate cancer patients before and after ADT with enzalutamide^74^, supports these findings. The upregulation of PGC-1α and OXPHOS following ADT suggests that ADT imposes a selection pressure on prostate cancer cells to rely more heavily on OXPHOS. This metabolic shift may facilitate progression toward the SCN state. Consequently, targeting the PGC-1α-OXPHOS axis during or immediately following ADT, rather than at the end-stage SCNCs, may offer an alternative therapeutic approach to curb SCN prostate cancer progression.

### Concluding remarks

Our study delineates the metabolic heterogeneity within SCNC subtypes, identifying distinct metabolic profiles and capacities for OXPHOS between the ASCL1 and POU2F3/ASCL2 subtypes. We demonstrate that PGC-1α not only distinguishes these subtypes but also drives metabolic reprogramming, which directly alters the cancer phenotype. These findings open new avenues for research and hold potential for guiding future therapies.

## MATERIALS AND METHODS

### PARCB + PGC-1α transformation

Donor prostate tissues were obtained in a de-identified manner and were therefore exempt from Institutional Review Board (IRB) approval. The PARCB transformation assay was performed as described previously^6^ with modifications to the viral constructs to include PGC-1α gain-and loss-of-function analyses. Briefly, donor prostate tissues were digested overnight and isolated cells were stained to identify the basal cells. FACS sorted basal cells were combined with PARCB oncogene lentiviruses at an MOI of 50 in Matrigel (Cat# 356234, Corning). The mixtures of basal cells, viral supernatant, and Matrigel were plated in 48-well plates as hanging drop cultures until Matrigel solidified into domes. 20,000 basal cells were used per Matrigel dome, in a final volume of 20-30 µl per well. After Matrigel solidification, organoids were cultured in prostate organoid media^78^ at 37°C and 5% CO2 for 10-14 days. Transduced organoids were harvested by dissociation of Matrigel with 1mg/mL Dispase (Cat# 17105041, Thermo Fisher Scientific) and washed three times with PBS to remove Dispase. Washed organoids were re-suspended in 10 μl of normal Matrigel and 10 μl Matrigel with high growth factors (Cat# 354248, Corning). The organoid-Matrigel mixtures were implanted subcutaneously in immunodeficient NOD.Cg-Prkdcscid Il2rgtm1Wjl/SzJ (NSG) mice^79^ to initiate tumor formation. Tumors were extracted before ulceration or reaching around 1 cm in diameter, whichever came first. NSG mice were obtained from the Jackson Laboratories and housed and bred under the care of the Division of Laboratory Animal Medicine at the University of California, Los Angeles (UCLA). All animal handling and subcutaneous injections were performed following the protocols approved by UCLA’s Animal Research Committee.

### Cell lines and culture media

The SCN prostate cancer line NCI-H660 (Cat# CRL-5813) was purchased from American Type Culture Collection (ATCC). The PARCB-P3-TP5, PARCB-P2-TP6, and PARCB-P7-TP6 cell lines^11^ and PARCB-1, PARCB-11^2^ cell lines were generated previously. The SCN Lung Cancer Cell lines CORL-311 (POU2F3 subtype), NCI-H209 (ASCL1 subtype) were purchased from ATCC. LNCaP (Cat# CRL-1740) and C4-2B (Cat# CRL-3315), cell lines were purchased from ATCC. SCN prostate and lung cancer cell lines were maintained in stem cell culture media (SCM): Advanced DMEM/F12 (Gibco CAT# 12634028), Glutamax (Gibco, CAT# 35050061), Pen/Strep, B27 (Gibco CAT# 17504044), 10ng/ml human EGF (Peprotech CAT# 100-47, and 10ng/ml human FGF-basic (Peprotech CAT# 100-18B). LNCaP and C4-2B cell lines were maintained in RPMI medium supplemented with 10% FBS, 100 U/mL penicillin and 100 μg/mL streptomycin, and 4 mmol/L GlutaMAX. All cell lines were routinely tested for Mycoplasma using a MycoAlert™ PLUS Mycoplasma Detection Kit (Cat# LT07-703, Lonza).

### Lentiviral vectors and high-titer lentivirus production

The following three vectors for PARCB transformation were described previously: mristoylated AKT1 (FU-myrAKT1-CGW), c-MYC and BCL2 (FU-cMYC-P2A-BCL2-CRW), dominant negative TP53 (R175H) and shRNA targeting of RB1 (FU-shRB1-TP53DN-CYW)^6^. For PGC-1α overexpression, a fourth vector was designed. First, an FUGW backbone was subcloned to replace the EGFP with EBFP2 using Gibson assembly (referred to as FUBW). Next, PGC-1α cDNA from pcDNA4 myc PGC-1 alpha (Addgene #10974) was cloned into the FUBW backbone driven by a ubiquitin promoter. An empty FUBW backbone without PGC-1α cDNA was used as a control. For PGC-1α inhibition during PARCB transformation, shRNA targeting PGC-1α was cloned into the myristoylated AKT1 vector (FU-myrAKT1-CGW), downstream of the H1 promoter. The following PGC-1α targeting sequence was used TATGACAGCTACGAGGAATAT. As a control, a sequence that targets no known mammalian genes was used (CAACAAGATGAAGAGCACCAA). For PGC-1α inhibition in cultured cell lines, the same shRNA sequences were cloned into a pLKO_005 backbone, which was subcloned to replace its puromycin resistance gene along with its promoter with CMV-EBFP2. High titer lentiviruses were made using a previously established method^162^.

### Transduction of cultured SCN cell lines

Cultured SCN cell lines, which grow in suspension as clusters, were mechanically dissociated using gentle pipetting with a P1000 pipette. 600,000 live cells were seeded in non tissue culture treated 12-well plates and transduced with high titer lentiviruses at an MOI of 2-5 in the presence of polybrene and ROCK inhibitor. The plates were subsequently centrifuged at 1,000 g for 90 min at RT, followed by overnight incubation. The following morning, 1 ml fresh SCM was added to each well and the cells were gently mixed with a P1000 to dislodge any adhered cells. 48-72 hours after transduction, BFP positive cells were sorted to enrich for transduced cells. After a 24-hour recovery period, cells were used for downstream analyses.

### PRNBSA-induced SCN prostate cancer transdifferentiation in vitro

PNRBSA transduction experiments were performed as described previously^77^. Cells were seeded in 6-well tissue culture plates at a density of 3 x 105 cells per mL in 3 mL of RPMI medium supplemented with 10% FBS, 100 U/mL penicillin and 100 μg/mL streptomycin, and 4 mmol/L GlutaMAX. Cells were transduced approximately 4-6 hours after seeding at a defined multiplicity of infection (MOI) of 4 for each lentivirus. 72 hours after transduction, cells were trypsinized, washed, and transferred to 100 mm tissue culture plates in 15 mL of neural stem cell media (N-SCM) consisting of Advanced DMEM/F12 medium supplemented with 1X serum-free B27, 10 ng/mL recombinant human bFGF, 10 ng/mL recombinant human EGF, 100 U/mL penicillin and 100 μg/mL streptomycin, and 4 mmol/L GlutaMAX. Media were replenished every 3-4 days. Cells were collected 14 days post-transduction for analysis.

### Cell proliferation assay

2,000 -10,000 cells per cell line in 3-6 replicates were seeded into black, opaque 96-well plates with a glass bottom. Cell content was measured on Days 1-6 using Cell Titer-Glo Luminescent Cell Viability Assay (Cat# G7570, Promega). Luminescence was measured at an integration time of 0.5 second per well. The number of cell doublings were calculated between the day of plating, and the termination of the assay, indicated on each figure panel. Because CTG uses an ATP-based readout, and since the mitochondrial inhibitors influence the amount of ATP in the cell, the internal GFP, driven by the AKT1 (FU-myrAKT1-CGW) vector was also quantified using microscopy and a plate reader. Consistent results were obtained with both techniques.

### Inhibition of PGC-1α and OXPHOS

For pharmacological inhibition of PGC-1α (with SR-18292) and OXPHOS (with IACS-010759 and IMT1B), 96-well plates were pre-filled with SCM media (described above) containing each of the inhibitors or DMSO as a control. Cell growth was quantified using the Cell Titer-Glo Luminescent Cell Viability Assay (Cat# G7570, Promega) as described below.

### RT-qPCR

Total RNA was isolated from cells using miRNeasy Mini Kit (Cat# 217004, Qiagen). cDNA was synthesized from 2 ug of total RNA using the SuperScript IV First-Strand Synthesis System (Cat# 18091050, Thermo Fisher). RT-qPCR was performed using SYBR Green PCR Master Mix (Cat# 4309155, Thermo Fisher). Amplification was carried out using the StepOne Real-Time PCR System (Cat# 4376357, Thermo Fisher) and analysis was performed using the StepOne Software v2.3. Relative quantification was determined using the Delta-Delta Ct Method. The following primer pair was used to amplify PGC-1α mRNA: CCTGCTCGGAGCTTCTCAAA CCCTTGGGGTCATTTGGTGA

### Tissue section, histology, and immunohistochemistry (IHC)

PARCB tumors were fixed by overnight nutation in 10% buffered formaldehyde (SF100-4) at 4℃ followed by three washes in 70% ethanol. Hematoxylin and eosin (H&E) staining was performed by UCLA’s Translation Pathology Core Laboratory (TPCL) using standard protocols. TPCL is a CAP/CLIA certified research facility in the UCLA Department of Pathology and Laboratory Medicine and a UCLA Jonsson Comprehensive Cancer Center Shared Facility. For immunohistochemistry, formalin-fixed, paraffin-embedded (FFPE) PARCB tumors were deparaffinized in xylene and rehydrated. Citrate buffer (pH=6.0) was used for antigen retrieval. The sections were incubated in citrate buffer and heated in a pressure cooker. 3% H2O2 in methanol was used to block endogenous peroxidase activity for 10 min at room temperature. The sections were blocked then incubated with primary antibodies overnight at 4°C. Anti-mouse/rabbit secondary antibodies were used to detect proteins of interest and DAB EqV substrate was used to visualize the staining. All components were included in the ImmPRESS Kit (MP-7801-15 and MP-7802-15, Vector Laboratories). The slides were then dehydrated and mounted with Xylene-based drying medium (Cat# 22-050-262, Fisher Scientific).

### Western blot

1 million viable cells were lysed on ice using RIPA lysis buffer (Cat#89900, Thermo Fisher). Protein concentrations were measured using the Pierce BCA Protein Assay Kit (Cat#: 23227, Thermo Scientific). Samples were electrophoresed on polyacrylamide gels (Cat# NW04120BOX, Thermo Fisher), transferred to PVDF membranes (Cat# IPVH00010, Millipore). Western blots were visualized using iBright CL1500 Imaging system (Cat#44114, Thermo Fisher).

### Antibodies

The following antibodies were used: PGC-1α ms monoclonal 4C1.3 (Sigma, ST1202), TOMM20 ms monoclonal (Santa Cruz Biotechnology, sc-17764), NCAM1/CD56 (Abcam, ab133345), and OXPHOS cocktail ms monoclonal (Abcam, ab110411).

### Bulk RNA sequencing and dataset collection

Tumors were dissociated into single cells, followed by cell sorting of quadruple colors (BFP, GFP, YFP, and RFP) by flow cytometry. Total RNA was extracted from the cell lysates using the Zymo Direct-zol RNA Miniprep Plus Kit (Cat. No: R2072). Quality check was performed on the Agilent 4200 Tapestation (Agilent Technology; cat. no. G2991BA). Libraries for RNA-Seq of PARCB with PGC-1α overexpression and control samples were prepared with KAPA Stranded mRNA-Seq Kit (Cat# KK8420, Roche). The workflow consists of mRNA enrichment and fragmentation. Sequencing was performed on Illumina Hiseq 3000 or NovaSeq 6000 for PE 2×150 run. Data quality check was done on Illumina SAV. Demultiplexing was performed with Illumina Bcl2fastq v2.19.1.403 software. Raw sequencing reads were processed through the UCSC TOIL RNA Sequencing pipeline for quality control, adapter trimming, sequence alignment, and expression quantification. Briefly, sequence adapters were trimmed using CutAdapt v1.9, sequences were then aligned to human reference genome GRCh38 using STAR v2.4.2a and gene expression quantification was performed using RSEM v1.2.25 with transcript annotations from GENCODE v23^83^.

The FASTQ files of the Park dataset^6^, Beltran dataset^33^, George dataset^32^, Rajan dataset^124^ (GSE48403), and Chen dataset were all processed through the TOIL pipeline with the same parameters to get RSEM expected counts. The TOIL-RSEM expected counts of TCGA pan cancer samples were obtained directly from UCSC Xena browser (https://xenabrowser.net/datapages) and RSEM read counts of pan-cancer cell lines from the Cancer Cell Line Encyclopedia (CCLE) were downloaded from DepMap Portal (DepMap Public 22Q1) (https://depmap.org/portal/download/all/). The RSEM counts of all combined datasets were upper quartile normalized with a pseuodocount of 1 and log2 transformed (referred to as log2 (UQN+1) counts) and filtered down to HUGO protein coding genes (http://www.genenames.org/) for the downstream analyses. SCLC subtypes^43^ and CRPC subtypes^54^ were previously defined.

Sequencing of high-grade serous ovarian carcinomas (HGSOC) was approved through UCLA Institutional Review Board (IRB) approved protocols (IRB #10-000727, IRB #20-001626). Tumor specimens were obtained from high-grade serous ovarian cancer patients who had given their informed consent. RNA was isolated from both solid tumors collected from primary and metastatic sites, and effusion samples such as ascites and pleural fluid. Dissociated or cryopreserved tumor fragments were used for isolation of RNA. These samples were collected at various disease time points: chemonaive, post adjuvant or neoadjuvant chemotherapy, and at disease recurrence. Presence of tumor cells was confirmed on histologic sections. RNA isolation, library preparation, and sequencing were conducted by the Technology Center for Genomics and Bioinformatics (TCGB) core facility at UCLA.

### Differential gene expression analysis and hierarchical clustering

Differential expression analysis was performed on raw RSEM expected count data of protein-coding genes using the R package DESeq2^98^.

### Gene set enrichment analysis (GSEA) and GSEA-squared

Differential gene expression analysis was first performed on raw RSEM expected count data of protein-coding genes on the Rajan 2014^124^ dataset between Pre-ADT and Post-ADT samples.

Using the fgsea R package (https://github.com/ctlab/fgsea), gene set enrichment analysis (GSEA) was then performed on the hallmark (H), canonical pathways (CP), and gene ontology (GO) gene sets from MSigDB^99^. Additional mitochondria related gene sets were also included from MitoCarta3.0. The ranked list of genes was generated using the signed log2 fold change and -log10 transformed p-values calculated by DESeq2^100^. GSEA results were then ranked by Normalized Enrichment Score (NES) to identify highly enriched gene sets. To investigate pathways broadly related to our highly enriched gene sets (e.g., oxidative phosphorylation and adhesion), GSEA-Squared analysis was performed. In short, gene sets were ranked by NES and marked for if they contained a key term from that category of terms in the pathway name. KS tests were then performed using ks.test.2 to assess the distribution of the category of terms^100^. The keys used were the following:

#### Oxidative Phosphorylation

OXIDATIVE PHOSPHORYLATION, ELECTRON TRANSPORT, MITOCHONDRIAL COMPLEX, NADH DEHYDROGENASE, MITOCHONDRIAL LARGE RIBOSOMAL; OXPHOS

#### Adhesion

ADHESION, ADHERENS

Differential gene expression analysis was also performed on raw RSEM expected count data of protein-coding genes on the Chen 2014^32^ dataset between HC6/ Class II/ ASCL1+ tumors and the rest of the samples in the dataset. Similar methods were used to perform gene set enrichment analysis on GO BP 2021 gene sets from MSigDB, except genes were ranked by their adjusted p-value.

### Principle component analyses

Unsupervised principal component analysis (PCA) was performed on log2 transformed upper quartile normalized RSEM expected count data. The prcomp function in R was run centered and unscaled. Data was projected onto the PCA framework by multiplying the rotation matrix by the projected data’s expression matrix.

### Small cell neuroendocrine (SCN) score

SCN score was calculated as previously described in Balanis et al^1^. Briefly, PC1 loadings were taken from a pan-cancer PCA which followed non-small cell cancers along their transdifferentiation trajectory towards an SCN phenotype. A higher SCN score indicates a more SCN-like phenotype, while a lower SCN score indicates a more normal, non-SCN-like phenotype.

### Motif analysis

ATAC sequencing data was obtained from Chen 2023 dataset^32^. The raw FASTQ files were processed using the published ENCODE ATAC-Seq Pipeline (https://github.com/ENCODE-DCC/atacseq-pipeline). The reads were trimmed and aligned to hg38 using bowtie2. Picard was used to de-duplicate reads, which were then filtered for high-quality paired reads using SAMtools. All peak calling was performed using MACS3. The optimal irreproducible discovery rate (IDR) thresholded peak output was used for all downstream analyses, with a threshold P value of 0.05. Other ENCODE3 parameters were enforced with the flag-encode3. Reads that mapped to mitochondrial genes or blacklisted regions, as defined by the ENCODE pipeline, were removed. The peak files were merged using bedtools merge to create a consensus set of peaks across all samples, and the number of reads in each peak was determined using bedtools multicov^101^. A variance stabilizing transformation was performed on peak counts using DESeq2^98^ and batch effects were removed using removeBatchEffect from limma^102^. DESeq2 was then used to variance stabilize transform read counts and determine hyper-and hypo-accessible peaks across HC5 and HC6 PARCB time course samples, using default parameters and without independent filtering or Cook’s cutoff. Peaks were called as hyper or hypo-accessible using abs(log2 fold change)>2 and adjusted p<0.05. Motif analysis was then run separately on hyper-or hypo-accessible peaks in the HC5 versus. HC6 comparison using HOMER^103^ with the flags-size 200 and-mask. Motifs were then ranked by their p-value for hyper-or hypo-accessible peak sets. Motifs specific to hyper or hypo accessible peaks were obtained by taking the rank difference of the motifs in the two lists.

### Single-cell RNA sequencing analyses

Single-cell expression levels of ASCL1, PGC-1α, ASCL2, and POU2F3 in PARCB time course tumors were extracted from our recently published study^11^. Gene expression was visualized using Uniform Manifold Approximation and Projection (UMAP) analysis as described previously^11^. The downstream quality control, as well as batch integration and correction of PARCB single cell RNA-seq data were performed as described previously^11^. Briefly, visualization of PARCB single cell RNA-seq data was performed using the Seurat (5.0.3) R package^104^. The top 30 principal components were used to perform UMAP analysis. Cell clustering was performed with FindNeighbors function with top 30 principal components and FindClusters function with resolution of 0.5.

### Seahorse respirometry

Respirometry experiments were performed on a Seahorse XF96 Extracellular Flux Analyzer (Agilent Technologies). PARCB tumor-derived cell lines were process and counted as described, washed into Seahorse Assay medium (Seahorse XF Base Medium supplemented with 2 mM L-glutamine, 1 mM pyruvate and 10 mM glucose) and immediately seeded into an XF96 microplate, pre-coated with PDL. Cells were plated at a density of 40,000 cells per well in a final volume of 175 µl. Prior to the start of the assay, the XF96 plate was placed in a 37 °C incubator without CO2 for 30 min. During the assay, the following compounds were injected: the mitochondrial ATP synthase inhibitor oligomycin (final concentration 2 μM); the mitochondrial uncoupler FCCP (final concentration 1 μM); and the complex I and III inhibitors rotenone (final concentration 2 μM); and the complex III inhibitor antimycin A (final concentration 2 μM). At the conclusion of the assay, the cells were fixed with 4% paraformaldehyde, stained with Hoechst, and cell number per well was determine based on nuclei number using an Operetta High-Content Imaging System (PerkinElmer). OCRs were normalized to cell number per well.

### microPET/CT imaging of tumor metabolism

Tumor-bearing mice were injected with 60 µCi of [^18^F]-FDG (PETNET Solutions) or [^18^F]-BnTP (prepared as previously described^105^ through tail vein intravenous injections. Following a 60-minute unconscious uptake of the imaging tracer, mice were anesthetized with 2% vaporized isoflurane, and microPET (energy window 350-650 keV, 10-min static scan) and microCT (voltage 80 kVp, current 150 μA, 720 projections, 200μm resolution, scan time 1 min) images were acquired on a GNEXT PET/CT scanner (Sofie Biosciences, Dulles, VA). The microPET images were reconstructed using a 3D-Ordered Subset Expectation Maximization (OSEM) algorithm (24 subsets and 3 iterations), with random, attenuation, and decay correction. The microCT images were reconstructed using a Modified Feldkamp Algorithm. Amide software was used to analyze co-registered microPET/CT images. Representative maximum-intensity-projection (MIP) images are shown.

### Chromatin immunoprocepticiation sequencing (ChIP-Seq)

To analyze the regulatory impact of ASCL1 on PGC-1α, we utilized Chromatin Immunoprecipitation Sequencing (ChIP-Seq) and CUT&RUN data from several studies, including Augustyn et al., 2014 (GSE61197), Borromeo et al., 2016 (GSE69398), Cejas et al., 2021 (GSE156290), and Nouruzi et al., 2022 (GSE183198), which were downloaded directly from the Gene Expression Omnibus (GEO). We converted data from bedGraph and wig formats to bigWig format using the UCSC bedGraphToBigWig and wigToBigWig tools to standardize data representation. To align all datasets to a consistent reference genome, we lifted data initially mapped to hg19 over to hg38 using the UCSC liftOver tool. For the Li et al., 2024 ChIP-Seq dataset^77^, we merged bigWig files from replicates at each timepoint using the UCSC bigWigMerge tool, aiming to consolidate data for comprehensive analysis. Visualization of the processed datasets was performed with the Integrative Genomics Viewer (IGV 2.17.1), ensuring uniform background noise levels across samples to facilitate accurate comparison.

## ACKNOWLEDGEMENTS

We thank the University of California, Los Angeles (UCLA) Tissue Procurement Core Laboratories for prostate tissue preparation and histological staining, and the UCLA Technology Center for Genomics and Bioinformatics for RNA sequencing. We thank Nicholas Bayley at UCLA for RNA-sequencing data processing. The microPET/CT imaging in this study was supported by the NIH Cancer Center Support Grant (2 P30 CA016042-44). The GNEXT PET/CT scanner was funded by NIH S10 Shared Instrumentation for Animal Research Grant (1 S10 OD026917-01A1). We thank Linsey Stiles and Katrina P. Montales of the UCLA Mitochondria and Metabolism Core for assistance with the XF96 Extracellular Flux Analyzer. The graphical abstract and Figure 2A were partially created with BioRender.com. Finally, we thank the following UCLA undergraduate research assistants who helped with wet lab experiments: Spencer Gaut, Alitzel Martinez, Michelle Garcia, and Kathy Rivera.

## AUTHOR CONTRIBUTIONS

<u>Conceptualization</u>: G.V., O.N.W.; <u>Methodology</u>: G.V., S.X., V.B., D.S., J.W.P., T.G., O.N.W.; <u>Formal Analysis</u>: G.V., C.C., J.F., T.H., W.T., K.S., M.H., F.N.E., L.W.; <u>Investigation</u>: G.V., D.C., V.B., F.N.E., G.A.B.; <u>Resources</u>: C.C., J.W.P., S.M.; <u>Writing-Original Draft</u>: G.V.; <u>Writing – review & editing</u>: G.V., E.A., O.N.W.; <u>Visualization</u>: G.V.; S.X., M.H.; <u>Supervision</u>: S.M., D.S., J.K.L, T.G., O.S., O.N.W.

