## Supplemental Figures and Legends for "PGC-1α drives small cell neuroendocrine cancer progression towards an ASCL1-expressing subtype with increased mitochondrial capacity"

**Figure S1.** Related to Figure 1

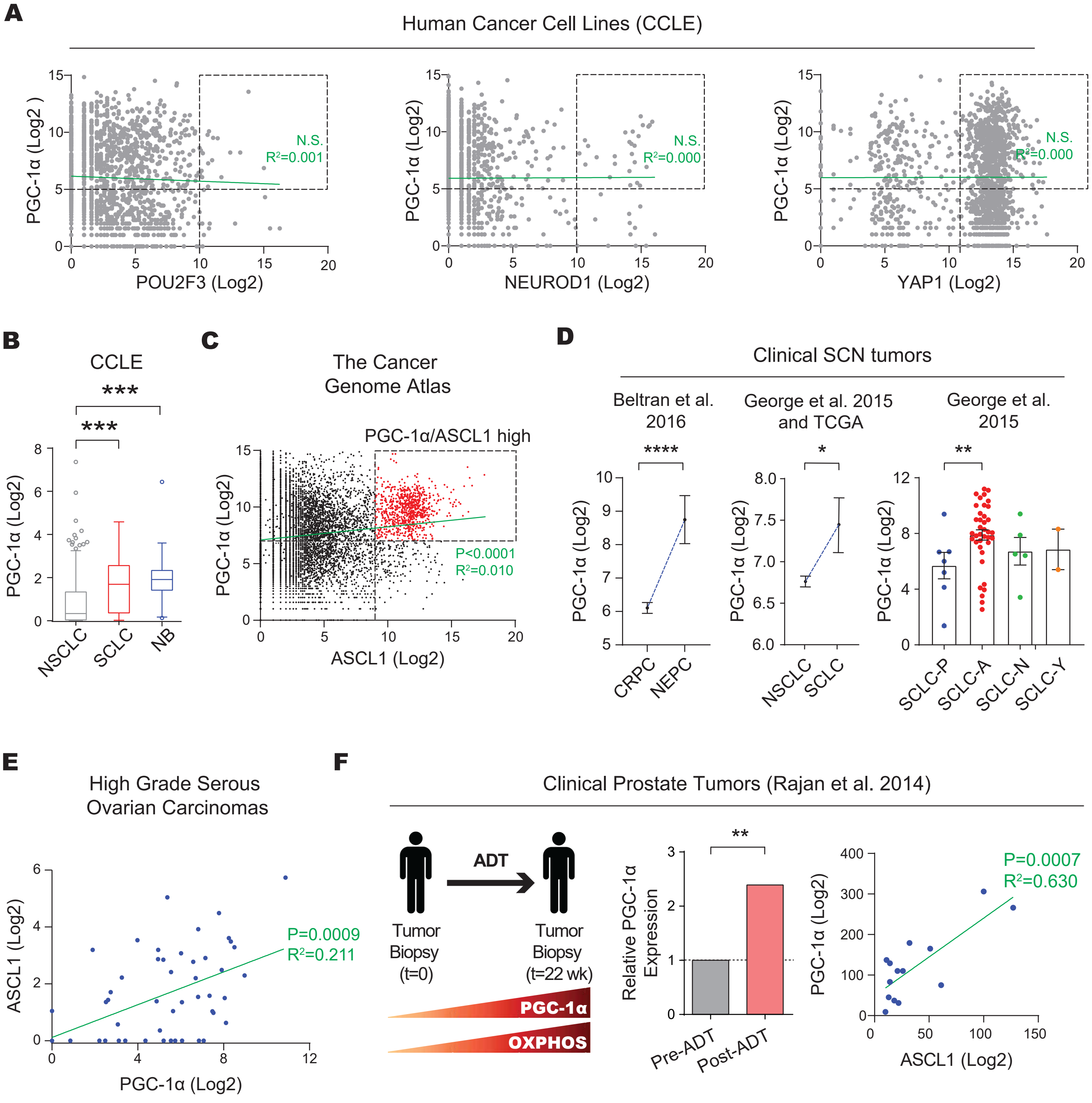

**Figure S1.** Related to Figure 1.

A. Related to Figure 1A. Gene expression analysis of PGC-1α versus the SCNC lineage markers POU2F3, NEUROD1, and YAP1 in all the human cancer cell lines from the cancer cell line encyclopedia (CCLE).

B. Related to Figure 1A. PGC-1α expression levels in non-small cell lung cancer (NSCLC), small cell lung cancer (SCLC), and neuroblastoma (NB) cell lines from the CCLE.

C. Co-expression analysis of PGC-1α and ASCL1 in all tumor samples from The Cancer Genome Atlas (TCGA).

D. PGC-1α expression levels in multiple SCNC datasets. SCLC subtypes are denoted as follows: P, POU2F3; A, ASCL1; N, NEUROD1; Y, YAP1. The datasets used are indicated.

E. Co-expression analysis of PGC-1α and ASCL1 in a cohort of patients with high-grade serous ovarian carcinomas (HGSOC).

F. Transcriptomic analyses in clinical prostate cancer tumors before and after androgen deprivation therapy (ADT) with enzalutamide. Left panel: a schematic overview. Middle panel: Comparison of PGC-1α expression levels pre- and post-ADT. Right panel: Co-expression analysis of PGC-1α and ASCL1 in (combined pre- and post-treatment analysis). See also Figure 1E.

The data for Figure S1B, S1D, and S1F are presented as mean ± SEM, with statistical significance denoted by *** ($p \leq 0.001$). The green lines represent the best-fit linear regression model, with the indicated $R^2$ and p-values. All Log2 values are Log2 UQN+1. For statistical tests used, see Material and Methods section

**Figure S2.** Related to Figure 2

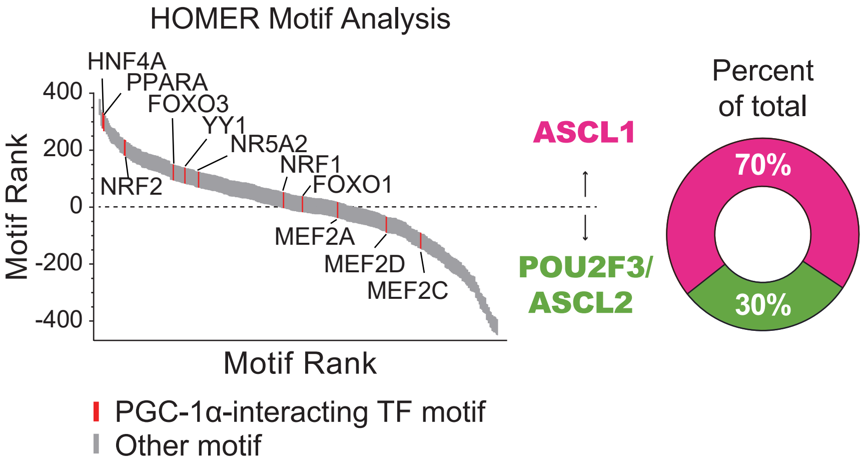

**Figure S2.** Related to Figure 2.

Hypergeometric Optimization of Motif EnRichment (HOMER) analysis across PARCB tumors from both POU2F3/ASCL2 and ASCL1 subtypes. PGC-1α interacting transcription factors were identified from the STRING database[108].

**Figure S3.** Related to Figure 2

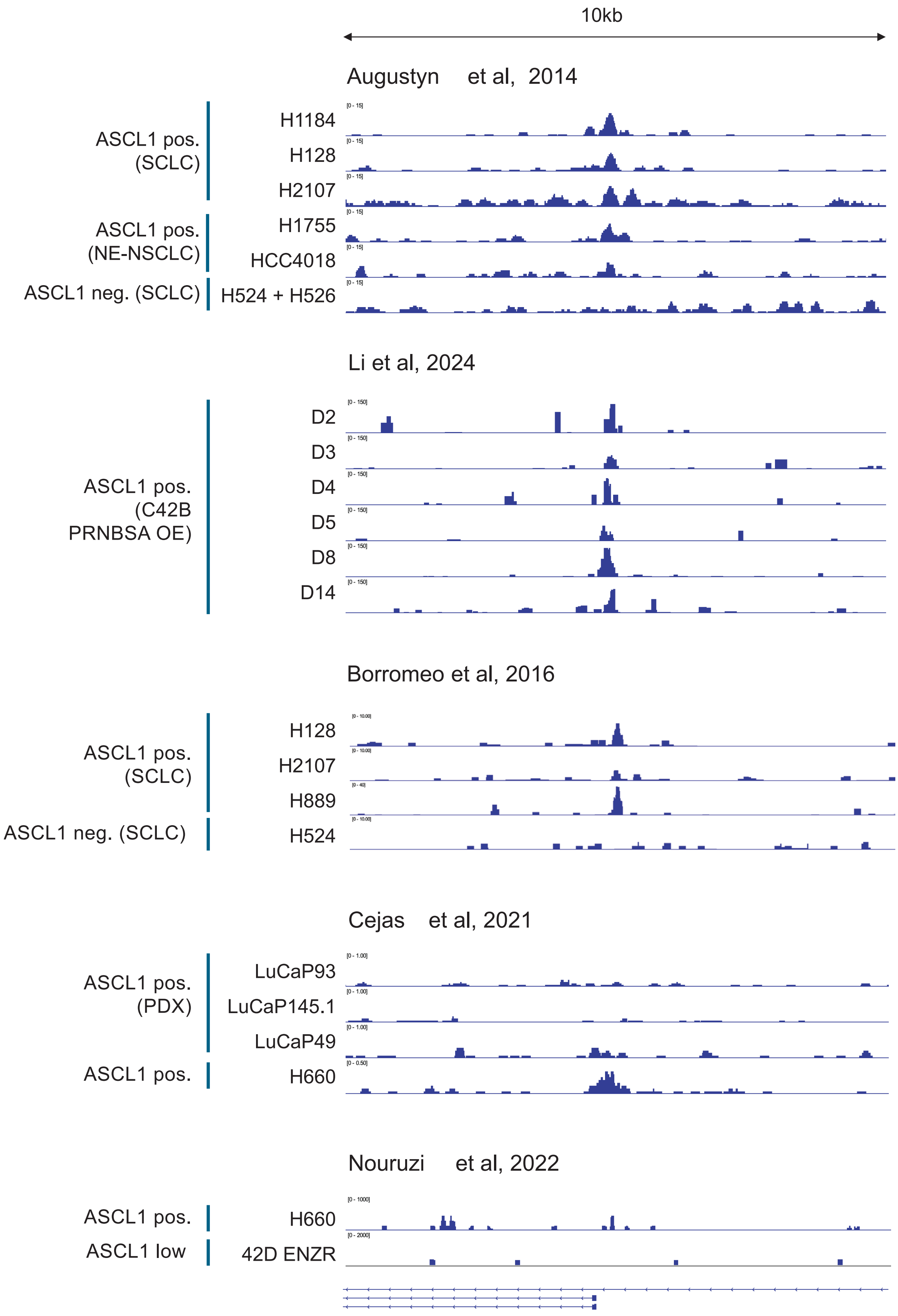


Promoter of PPARGC1A
Exon 1b, transcript variant X2 (XM_005248132.1)

**Figure S3.** Related to Figure 2

ASCL1 ChiP-Seq analyses showing the alternative promoter of PGC-1α corresponding to variant X2 (XM_005248132.1) encoded by exon1b. The following datasets were used: SCN lung cancer cell lines[75,76], SCN prostate cancer cell lines[63,77], and LuCaP prostate cancer patient-derived xenografts (PDXs)[10].

**Figure S4.** Related to Figure 2

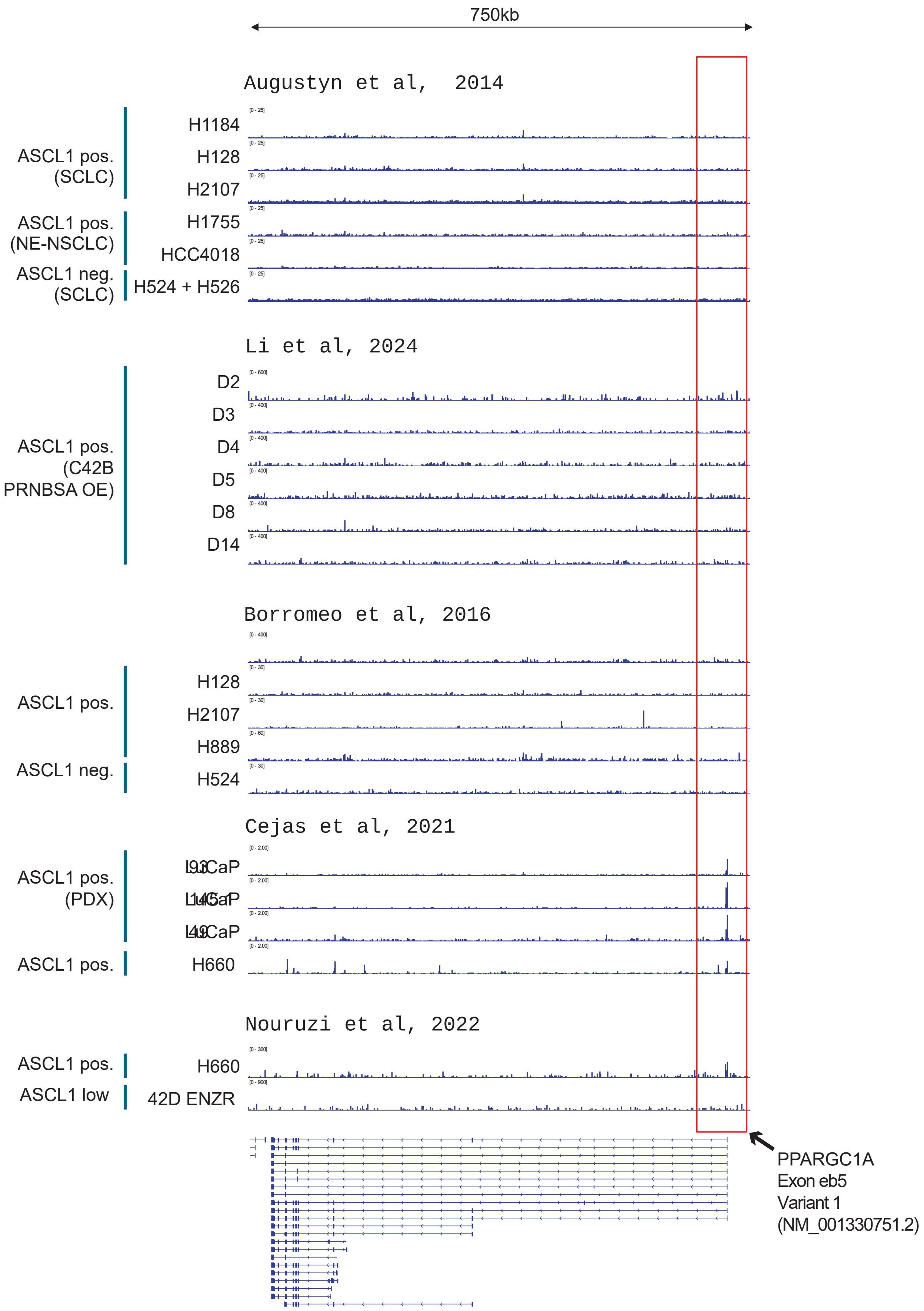

**Figure S4.** Related to Figure 2

ASCL1 ChiP-Seq analyses highlighting variant 1 of PGC-1α (NM_001330751.2) (red box). The following datasets were used: SCN lung cancer cell lines[75,76], SCN prostate cancer cell lines[63,77], and LuCaP prostate cancer patient-derived xenografts (PDXs)[10].

**Figure S5.** Rrelated to Figure 2

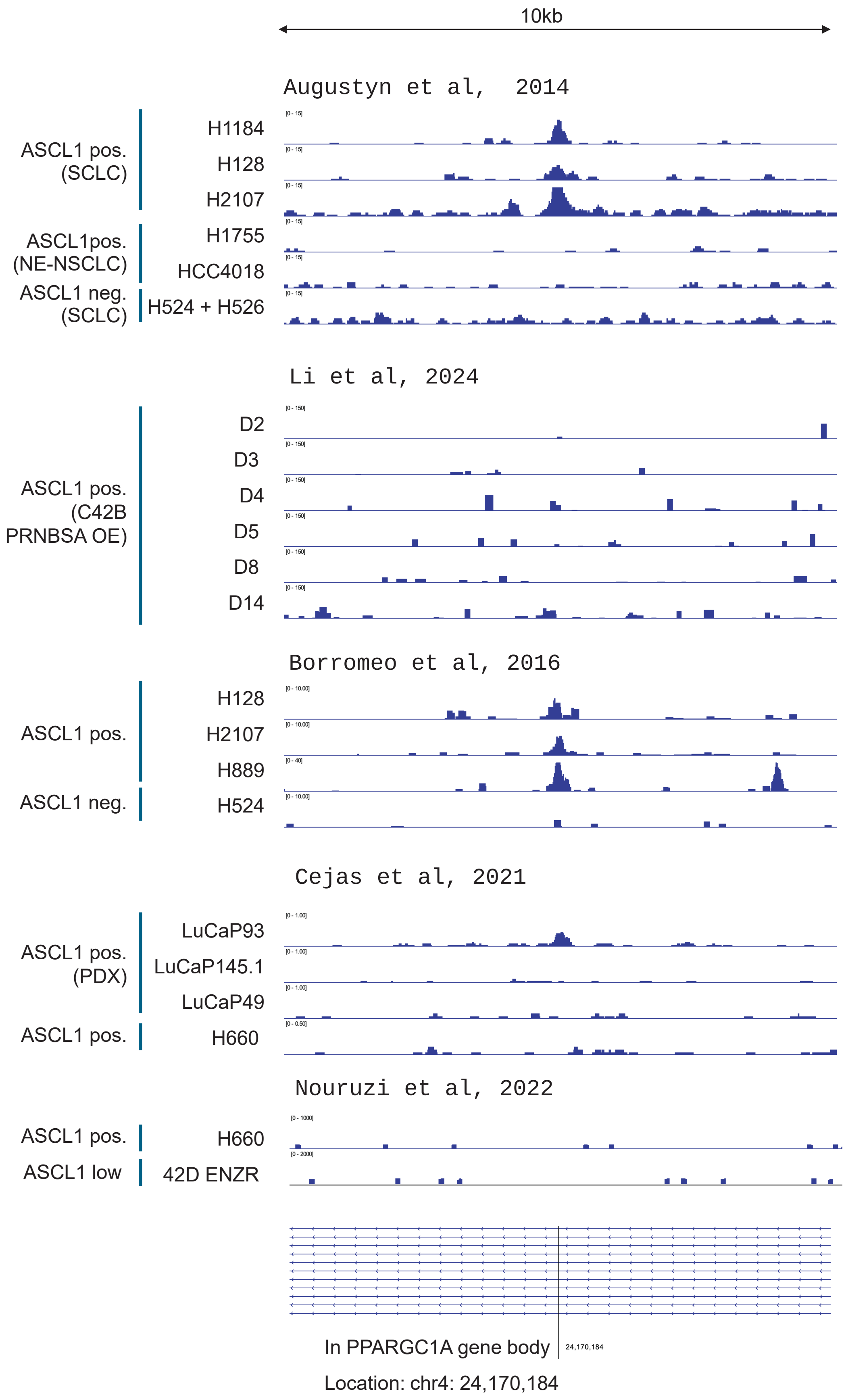

**Figure S5.** Related to Figure 2

ASCL1 ChiP-Seq analyses indicating a region within the PGC-1α gene body. The following datasets were used: SCN lung cancer cell lines[75,76], SCN prostate cancer cell lines[63,77], and LuCaP prostate cancer patient-derived xenografts (PDXs)[10].

**Figure S6.** Related to Figure 2

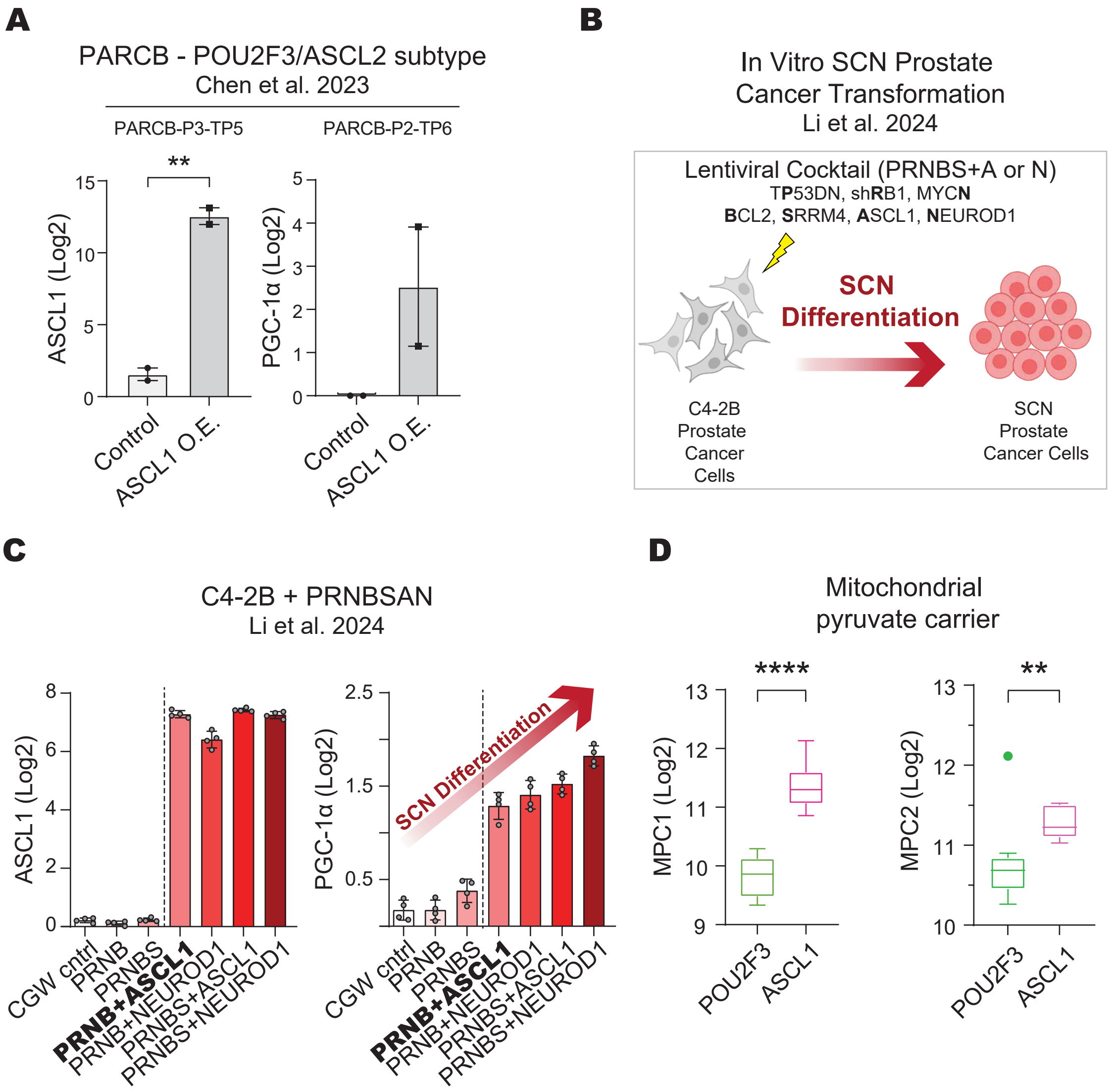

**Figure S6.** Related to Figure 2

A. Expression levels of ASCL1 (left panel) and PGC-1α (right panel) after acute overexpression of ASCL1 in cell lines derived from the PARCB POU2F3/ASCL2 tumor subtype. Data are mined from Chen et al 2023[11].

B. Schematic illustrating transformation of C42B cells to SCN prostate cancer using the PRNBSAN oncogenes and transcription factors (dominant-negative TP53, shRB1, MYCN, BCL2, SRRM4, ASCL1, and NEUROD1).

C. Expression analysis of ASCL1 (left panel) and PGC-1α (right panel) from C4-2B cells transduced with PRNBSAN. Log2 values are Log2 FPKM+1. The arrow indicates increased SCN differentiation as observed Li et at. 2024[77].

D. Expression of the mitochondrial pyruvate carrier in ASCL1 and POU2F3/ASCL2 PARCB tumor subtypes.

The data for Figure S6A and S6D are presented as mean ± SEM, with statistical significance denoted by ** ($p \leq 0.01$). Log2 values are Log2 UQN+1 unless otherwise noted. For statistical tests used, see Material and Methods section.

**Figure S7.** Related to Figure 3

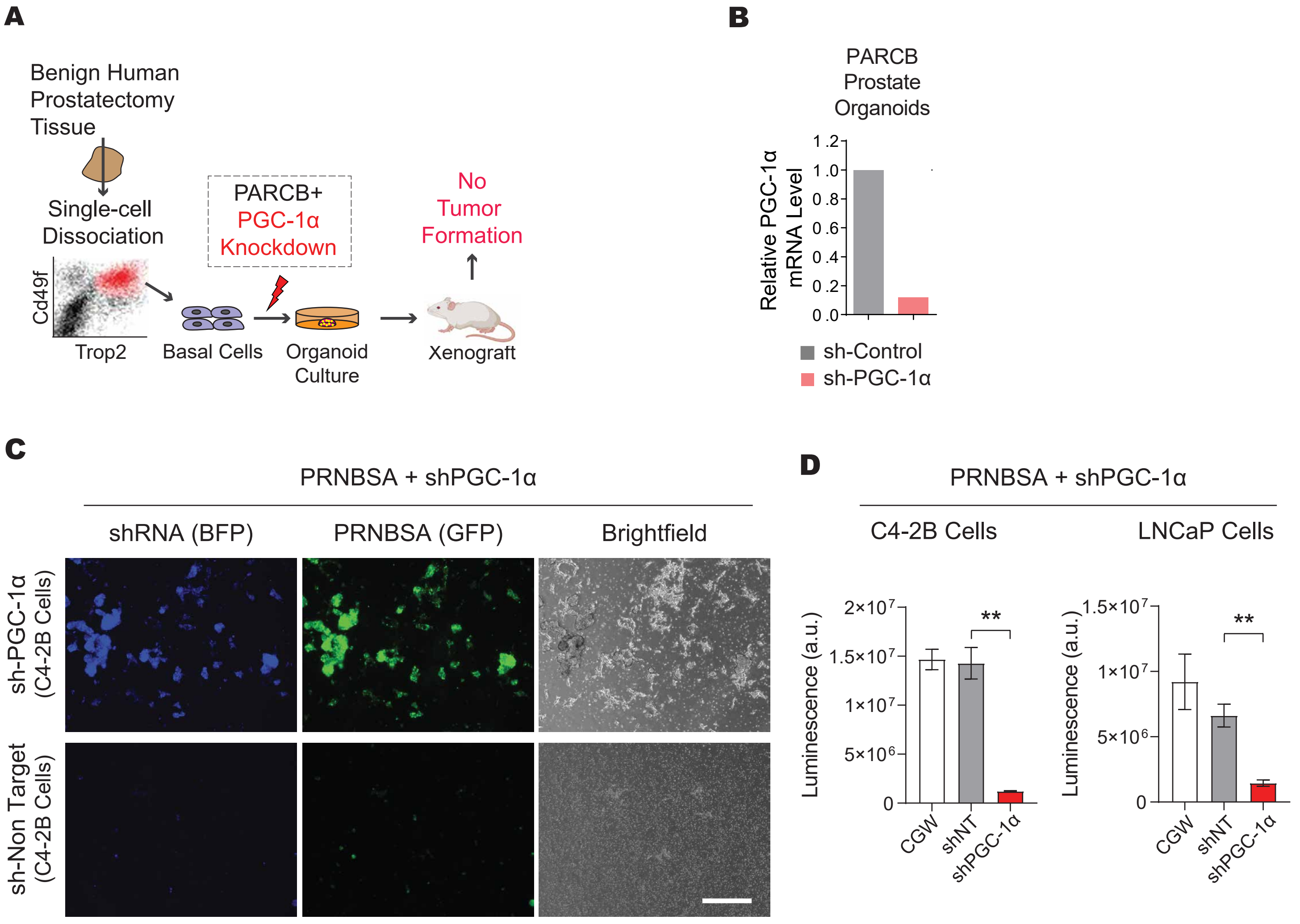

**Figure S7.** Related to Figure 3

A. Schematic illustrating PARCB prostate transformation with PGC-1α knockdown.
B. PGC-1α mRNA levels using RT-qPCR in PARCB-transduced prostate organoids.
C. PGC-1α inhibition during PRNBSA-mediated SCN prostate cancer differentiation in vitro. Representative images are shown in C4-2B cells.
D. Cell viability 14 days post transduction PRNBSA and indicated shRNAs. Quantification is shown from both C4-2B (left panel) and LNCaP cells (right panel).

**Figure S8.** Related to Figure 5

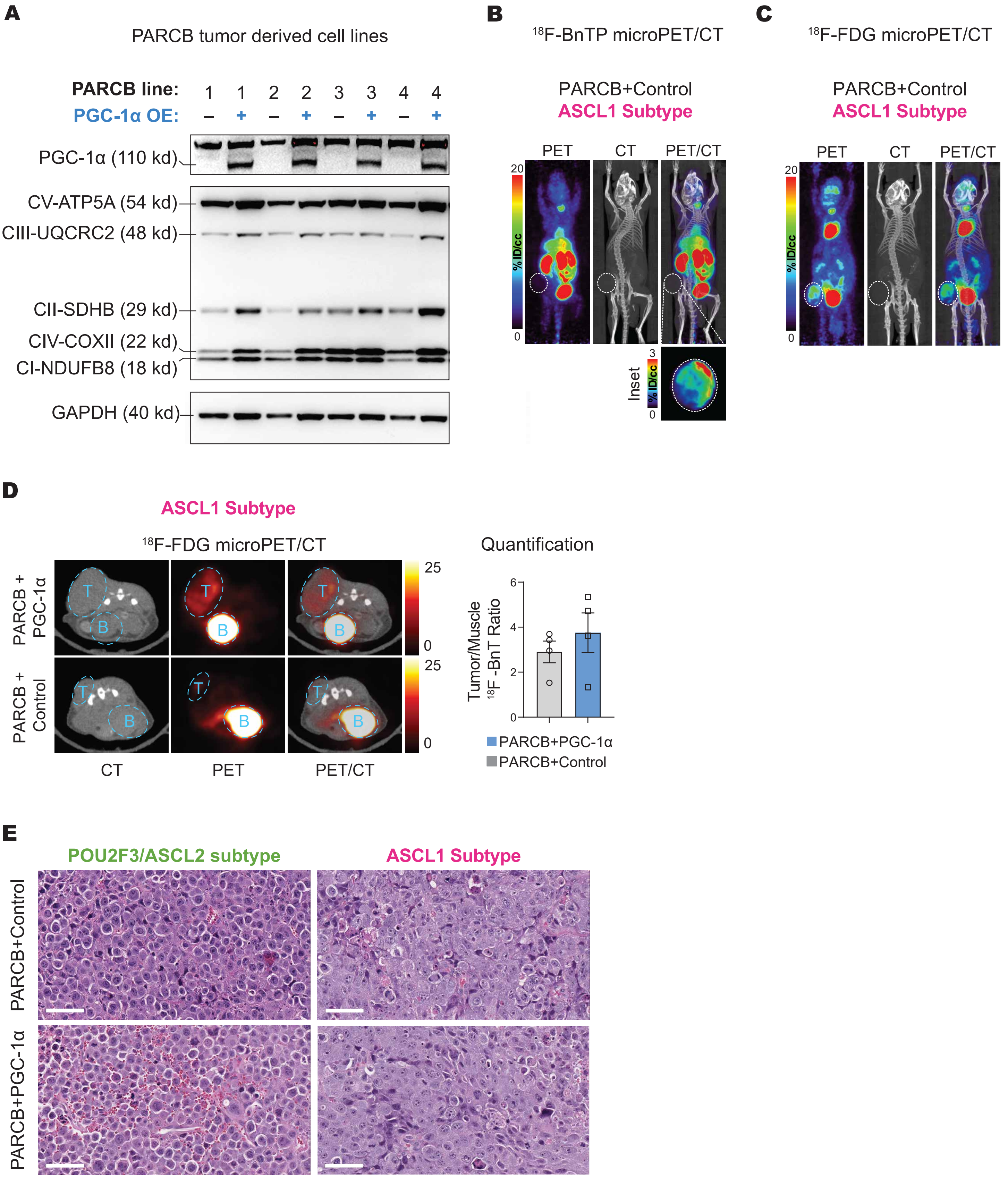

**Figure S8.** Related to Figure 5

A. Western blot analysis to evaluate the effect of PGC-1α overexpression on the expression levels of all five respiratory chain complexes across a panel of four PARCB tumor-derived cell lines. The PARCB cell lines were generated previously[2]

B. Overlay of in vivo microPET/CT scanning of a mouse with a subcutaneous ASCL1 PARCB tumor imaged with $^{18}$F-BnTP indicting mitochondrial membrane potential.

C. Overlay of in vivo micro PET and computed tomography scanning of a mouse with a subcutaneous ASCL1 PARCB tumor imaged with $^{18}$F-FDG, indicting glucose uptake.

D. Left panel: Representative $^{18}$F-FDG transverse PET-CT images of mice with subcutaneous tumor implantation. Uptake of PET probe was measured as the maximum percentage of injected dose per cubic centimeter (ID%/cc). Tumors are labeled “T”, and bladders are labeled “B”. Right panel: quantification of $^{18}$F-FDG uptake in the indicated groups. Values are normalized to PET signal from adjacent skeletal muscle.

E. H&E staining in tumor sections from POU2F3/ASCL2 and ASCL1 PARCB tumors with PGC-1α overexpression and control. These tumors were generated from terminally differentiated PARCB cell lines re-injected into mice for tumor growth, as described in Figure 5. Note the increased vasculature of POU2F3/ASCL2 tumors upon PGC-1α overexpression. Scale bars are 60 μm.

**Figure S9.** Related to Figure 6

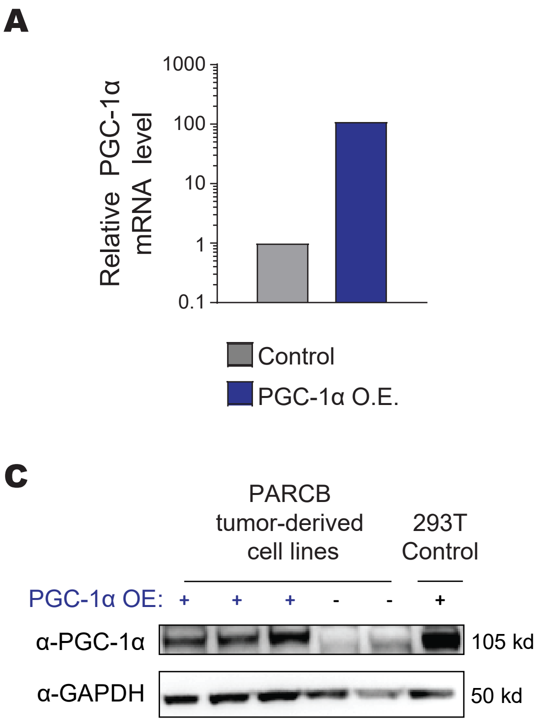


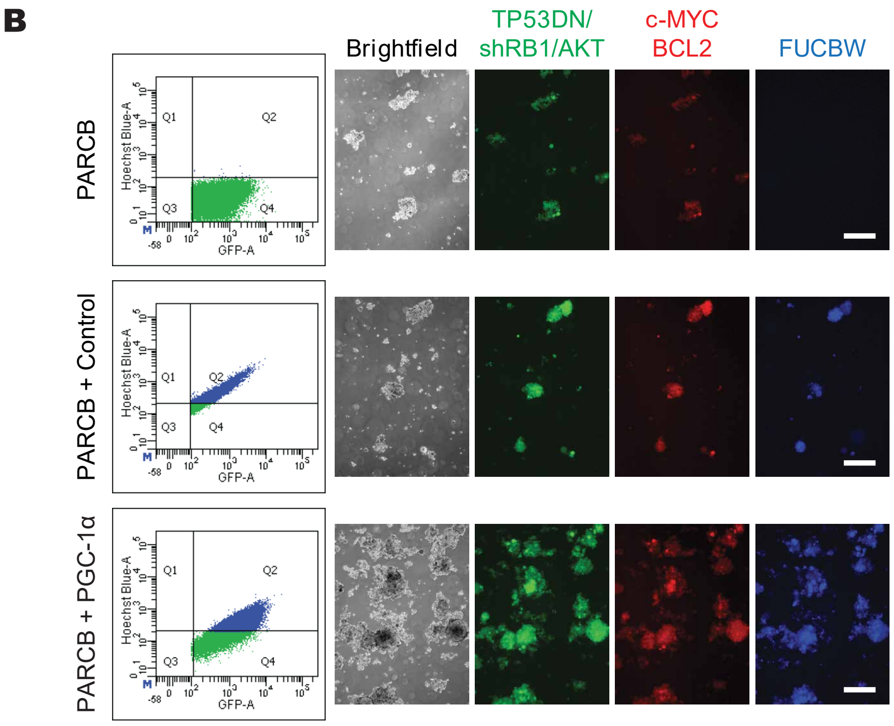

**Figure S9.** Related to Figure 6

A. PGC-1α mRNA levels using RT-qPCR in PARCB organoids with PGC-1α overexpression versus control. Organoids were harvested just before xenografting.

B. Flow cytometry and fluorescence microscopy analysis of PARCB tumors with PGC-1α overexpression and control. The PGC-1α overexpression and control constructs include a blue fluorescence protein encoded by EBFP2. See Materials and Methods section for more details.

Western blot analysis of PGC-1α protein levels in PARCB tumors with PGC-1α overexpression versus control. HEK 293T cells overexpressing PGC-1α were included as an additional control.
